# Age-related Macular Degeneration patient deep phenotyping and whole genome sequencing analysis identifies coding variants linking small low-luminance visual deficit to fat storage defects

**DOI:** 10.1101/2022.06.22.497149

**Authors:** Sehyun Kim, Amy Stockwell, Han Qin, Simon S. Gao, Meredith Sagolla, Ivaylo Stoilov, Arthur Wuster, Phillip Lai, Brian L. Yaspan, Marion Jeanne

## Abstract

**Background:** The basis of Age-related macular degeneration (AMD) genetic risk has been well documented; however, few studies have looked at genetic biomarkers of disease progression or treatment response within advanced AMD patients. Here we report the first genome-wide analysis of genetic determinants of low-luminance vision deficit (LLD), which is seen as predictive of visual acuity loss and anti-VEGF treatment response in neovascular AMD patients.

**Methods:** AMD patients were separated into small- and large-LLD groups for comparison and whole genome sequencing was performed. Genetic determinants of LLD were assessed by common and rare variant genetic analysis. Follow-up functional analysis of rare coding variants identified by the burden test was then performed *in vitro*.

**Results:** We identified four coding variants in the *CIDEC* gene. These rare variants were only present in patients with a small LLD, which has been previously shown to indicate better prognosis and better treatment response. Our *in vitro* functional characterization of these *CIDEC* alleles revealed that all decrease the binding affinity between CIDEC and the lipid droplet fusion effectors PLIN1, RAB8A and AS160. The rare *CIDEC* alleles all cause a hypomorphic defect in lipid droplet fusion and enlargement, resulting in a decreased fat storage capability in adipocytes.

**Conclusions:** As we did not detect CIDEC expression in the ocular tissue affected by AMD, our results suggest that the *CIDEC* variants do not play a direct role in the eye and influence low-luminance vision deficit via an indirect and systemic effect related to fat storage capacity.

**Funding:** No external funding was received for this work.

## Introduction

Age-related macular degeneration (AMD) accounts for nearly 10% of blindness worldwide, and is the leading cause of blindness in developed countries^1^. AMD is a progressive retinal disease characterized by the accumulation of extracellular deposits called drusen, underneath the retina in the early stages of the disease, followed by either atrophy of the macula in the advanced dry form of AMD called Geographic Atrophy (GA), and/or growth of pathogenic blood vessels into the retina in the wet form of AMD called neovascular AMD. Both GA and neovascular AMD are clinical end-stages forms of AMD and lead to progressive and severe vision loss. There is currently no approved treatment for GA and despite anti-Vascular Endothelial Growth Factor (VEGF) intraocular injections having revolutionized the treatment of neovascular AMD, they are not curative and patient response is heterogeneous^2^.

Although the pathophysiology of AMD is still not completely understood, there is a well-established genetic component to disease risk. Concordance rates between mono-zygotic twins are significantly higher than di-zygotic twins ^3–5^. Both population-based and familial studies have found evidence of sibling correlations, and estimate that genetic factors can account for between 50% and 70% of the total variability in disease risk ^6; 7^. Furthermore, it is estimated that genetic risk factors account for up to 71% of variation in the severity of disease^8^. Genome-wide association studies (GWAS) of AMD disease risk have greatly expanded our knowledge around the disease and especially its biology, with the most recent study involving over 16,000 AMD patients and 17,000 controls finding 52 independently associated variants ^9^. Major risk loci identified include complement genes (e.g. *CFH, CFI, C3, C9*) and the *ARMS2/HTRA1* locus. However, there are several other pathways identified including genes involved in lipid metabolism (e.g. *LIPC, CETP*) and extracellular matrix remodeling (e.g. *TIMP3, MMP9*).

While the basis of genetic risk of AMD is well characterized, other facets of the disease are not. Predictive or prognostic biomarkers, either clinical or genetic, for disease progression or treatment response are not as well understood. It is known that subjects with AMD have difficulty seeing in dimly lit environments ^10^. As such, the reduction in visual acuity under suboptimal illumination known as low-luminance deficit (LLD) has been evaluated in AMD patients and is seen to be predictive of both the development of GA with subsequent visual acuity loss and response to anti-VEGF treatment in neovascular AMD patients ^11; 12^.

Here we report the first genome-wide investigation into genetic determinants for low-luminance dysfunction in neovascular AMD utilizing patient data from the HARBOR clinical trial^13^. The HARBOR trial was a dosing study which sought to determine the efficacy and safety of 2.0 mg and 0.5 mg doses of ranibizumab (anti-VEGF antibody) in treatment naive patients with choroidal neovascularization (CNV) secondary to AMD^13; 14^. This study enrolled 1098 patients and followed them for one year. All dosing groups demonstrated clinically meaningful visual improvement. Multiple clinical datapoints were collected at baseline, including LLD. We separated the HARBOR patients into two groups for comparison, those with the largest LLD differential (biggest drop in vision under low-luminance, quartile 4 = Q4) and those with the smallest LLD differential before ranibizumab treatment (quartile 1 = Q1). We selected phenotypic extremities instead of the whole patient population for two main reasons; (1) the data looking at the effect of baseline LLD on anti-VEGF treatment response showed the largest difference between Q1 and Q4 patients^12^ and (2) it has been suggested as a way to increase power in genetic studies^15^. Because the genetic underpinnings of LLD differential has not been fully explored, we entered the study with the goal of identifying genetic factors involved in LLD using common and rare variation assayed via whole genome sequencing (WGS) with functional follow-up of biologically interesting hits. For functional characterization, we then selected from the top hits the *CIDEC* gene as a compelling candidate gene with reported function related to lipid metabolism, a pathway identified in previous AMD genetic analyses^9^.

## Results

### A genome wide burden test identifies rare genetic variants in the *CIDEC* gene that are enriched in AMD patients with small low-luminance deficit

For our study, we subset the HARBOR ranibizumab dosing study population as previously described for baseline low-luminance deficit (LLD) ^12^. All patients in the HARBOR trial had neovascular AMD. This subsetting resulted in 275 patients in our Q1 group, and 241 patients in Q4 (**Figure 1A**). Detailed population characteristics are seen in **Table 1**. We compared LLD quartiles 1 (Q1) and 4 (Q4) for this analysis, with the goal of maximizing the phenotypic difference as seen in the previous anti-VEGF treatment response study ^12^. Patients in Q1 (smallest low-luminance deficit) were seen to have better outcome on anti-VEGF therapy, and slower visual acuity loss in GA patients than patients in Q4 (large low-luminance deficit) ^11; 12^. In our study population, patients in Q1 were more likely to have lower baseline visual acuity, smaller baseline CNV leakage area, thinner sub-retinal fluid and a thinner choroid, but did not significantly differ by age or sex (**Table 1**). We coded Q1 as the “cases” and Q4 as the “controls”, so subsequently an odds ratio (OR)>1 indicates the minor allele was enriched in Q1 and an OR <1 indicates the minor allele was enriched in Q4.

**Figure 1:**
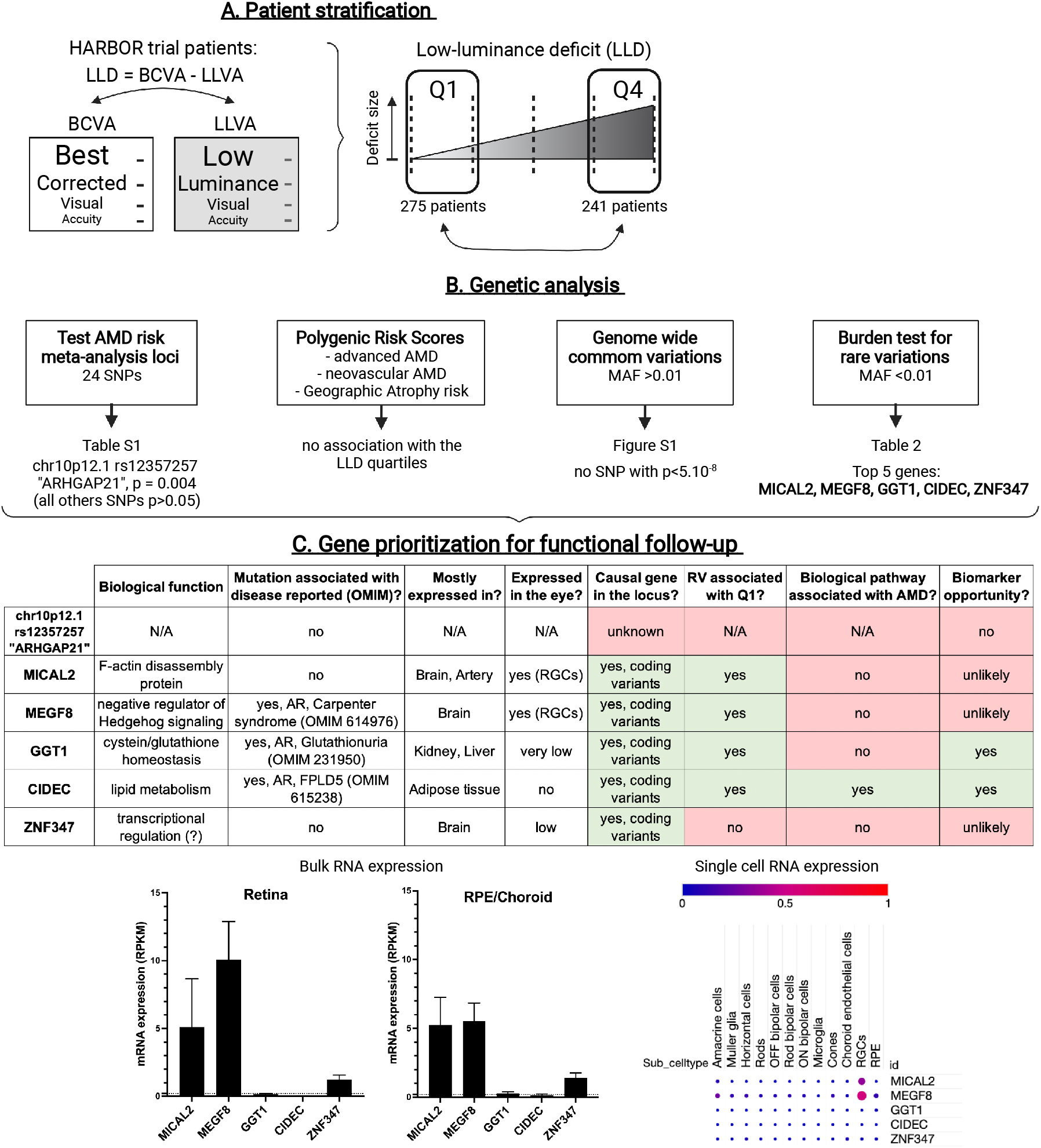
Overview of the patient stratification, the lines of genetic investigation performed and the strategy used to prioritize genes for functional follow-up. (A). HARBOR patients were separated at baseline into two groups based on the size of their Low-Luminance Deficit (LLD): patients in quartile 1 (Q1) had the smallest drop in vision under low-luminance and patients in quartile 4 (Q4) had the biggest deficit. (B). Lines of genetic investigation and top-line results. (C) For functional analysis follow-up, top genetic hits were prioritized based on different criteria such as being the causal gene at the locus (presence of coding variants), the rave variants (RV) identified being enriched in Q1 patients, the gene playing a role in a biological pathway associated with AMD pathophysiology, and providing a potential biomarker opportunity. For the top hits, gene expression in human retina or RPE/choroid (bulk RNA sequencing, data from Orozco et al.^34^) and in different human ocular cell types (single cell RNA sequencing, data from Gautam et al.^35^) were also analyzed. AR: autosomal recessive; FPLD5: Familial Partial Lipodystrophy type 5. RGCs: Retinal Ganglion Cells. RPE: Retinal Pigment Epithelium.

**Table 1.**
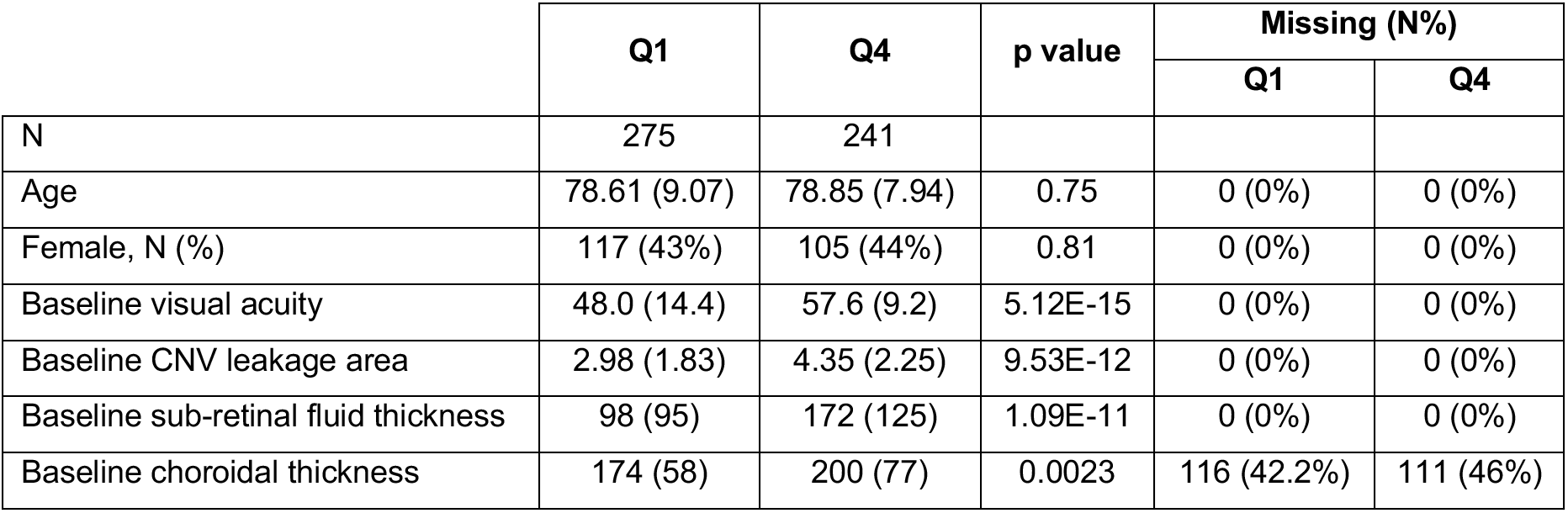
Quartile Q1 and quartile Q4 AMD patient demographic comparison.

|  | Q1 | Q4 | p value | Missing (N%) |  |
| --- | --- | --- | --- | --- | --- |
|  |  |  |  | Q1 | Q4 |
| N | 275 | 241 |  |  |  |
| Age | 78.61 (9.07) | 78.85 (7.94) | 0.75 | 0 (0%) | 0 (0%) |
| Female, N (%) | 117 (43%) | 105 (44%) | 0.81 | 0 (0%) | 0 (0%) |
| Baseline visual acuity | 48.0 (14.4) | 57.6 (9.2) | 5.12E-15 | 0 (0%) | 0 (0%) |
| Baseline CNV leakage area | 2.98 (1.83) | 4.35 (2.25) | 9.53E-12 | 0 (0%) | 0 (0%) |
| Baseline sub-retinal fluid thickness | 98 (95) | 172 (125) | 1.09E-11 | 0 (0%) | 0 (0%) |
| Baseline choroidal thickness | 174 (58) | 200 (77) | 0.0023 | 116 (42.2%) | 111 (46%) |

The lines of genetic investigation are outlined in **Figure 1B**. We first investigated the loci identified in a recent AMD risk meta-analysis from the International AMD Genetics Consortium (IAMDGC) (**Table S1**) ^8^. After quality control procedures, 24 single-nucleotide polymorphisms (SNPs) identified in the IAMDGC study were available for analysis. No locus retained statistical significance after multiple testing. Two loci had P<0.1, (1) *ARHGAP21*, rs12357257, (odds ratio (OR) = 0.63, p=0.004) and (2) *LIPC,* rs2043085, (OR = 1.29, P=0.10). We also constructed polygenic risk scores (PRS) for 1) advanced AMD risk 2) neovascular AMD risk and 3) geographic atrophy risk from the same IAMDGC consortium analysis. We did not find any of these PRS to be associated with our LLD population. Next, we examined common variation throughout the genome (SNPs with a minor allele frequency (MAF) > 0.01). There were no SNP which met the genome-wide significance level of p<5×10^−8^ (**Figure S1**).

We then evaluated rare variation (SNPs with a MAF<0.01) in the form of a burden test. We included exonic SNPs predicted to have a moderate (e.g. amino acid changing) to high (e.g. stop codon gain or loss) impact on the final protein sequence. No loci identified in the recent GWAS meta-analysis were significantly associated in our burden test (all p>0.05). No gene burden test passed a Bonferroni multiple testing cutoff for the number of genes in the genome tested. The top hits for this analysis are presented in **Table 2**.

**Table 2.**
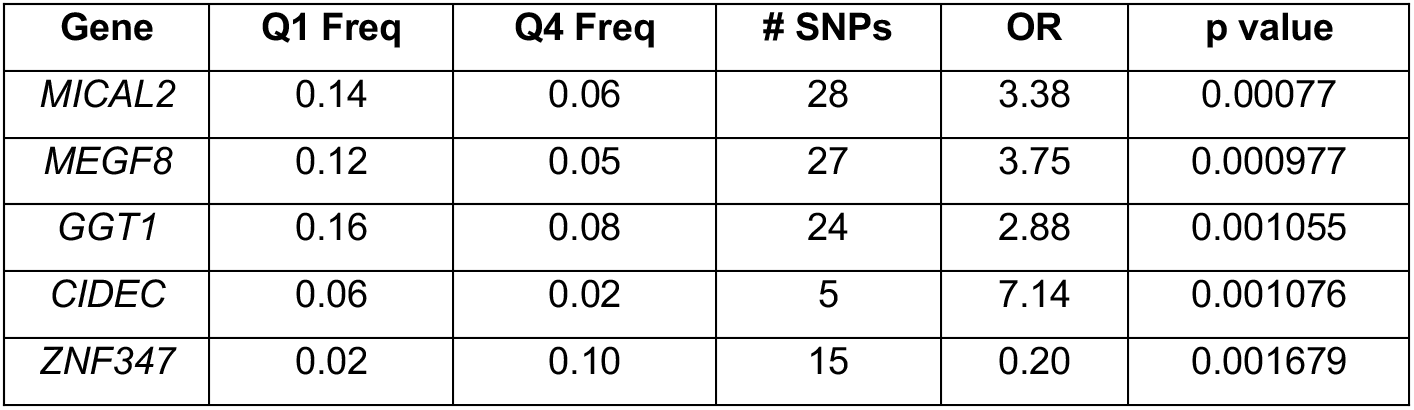
Results from rare variant burden test comparing quartile Q1 and quartile Q4 AMD patients. OR = odds ratio.

For functional analysis follow-up, we prioritized the top genetic hits identified by our common and rare variants analysis using different criteria (**Figure 1C**). We decided to select *CIDEC* for a thorough wet lab analysis as it was the probable causal gene at the identified locus (presence of coding variants), the rare variants identified were enriched in Q1 patients (i.e. associated with better outcome). Furthermore, CIDEC is involved in a biological pathway already associated with AMD (i.e. lipid metabolism) and since CIDEC expression is broad in the human body (adipose tissue), it provides a potential biomarker opportunity^16^, which is usually not the case when the gene expression is restricted to the neuroretina.

The *CIDEC* gene encodes the CIDEC protein (NP_001365420.1; OMIM: 612120), a member of the <u>C</u>ell-death-<u>I</u>nducing <u>D</u>NA fragmentation factor (DFF)45-like <u>E</u>ffector (CIDE) family. As this is the first report of CIDEC affecting AMD pathology, we looked for evidence of *CIDEC* rare variant involvement in the UK Biobank sequencing data via the GENEBASS portal (v0.7.8alpha)^17^. We did not see evidence of a strong phenotype associated with rare variants in *CIDEC* with regards to any distinct ocular phenotype, with “eye problems/disorders” being the top ocular phenotype in the pLoF analysis (**Table S2**; P=0.005).

The *CIDEC* rare alleles found in our analysis were found in 6% of Q1 patients and spread over multiple exons. In the Q4 patients, rare alleles were found in 2% of individuals and they coalesced to one exon seen only in RefSeq transcript NM_001199551 (**Figure 2A**). We sought to quantify the percentage of transcripts expressed that are NM_001199551 in the GTEx database for adipose tissue and blood ^18^. In both sample types, percent expression of NM_001199551 was 0.5% of all *CIDEC* transcripts (**Figure S2 – adipose pictured, blood similar**). In conclusion, if restricting the analysis in *CIDEC* to exons contained in transcripts that are more widely expressed we found that *CIDEC* rare variation was exclusive to the Q1 AMD patients (N=12). The four SNPs identified in Q1 patients were rs150971509 c.139G>A [p.Val47Ile], rs79419480 c.181T>C [p.Tyr61His], rs145323356 c.481G>A [p.Val161Met] and rs52790883 c.660G>T [p.Gln220His] (subsequently referred to as V47I, Y61H, V161M and Q220H respectively) (**Figure 2B**). We used the software PolyPhen-2 (Polymorphism Phenotyping v2) to perform *in silico* prediction of the possible impact of these four amino acid substitutions on CIDEC stability or function ^19^. The V47I substitution was predicted as probably damaging, the V161M and Q220H substitutions were predicted as possibly damaging and only the Y61H substitution was predicted to be benign. Since no structural data was available for the full CIDEC protein, these predictions were based solely on evolutionary comparisons. Thus, we decided to include all four rare variants identified in our Q1 AMD CIDEC patients in our experimental follow-up.

**Figure 2.**
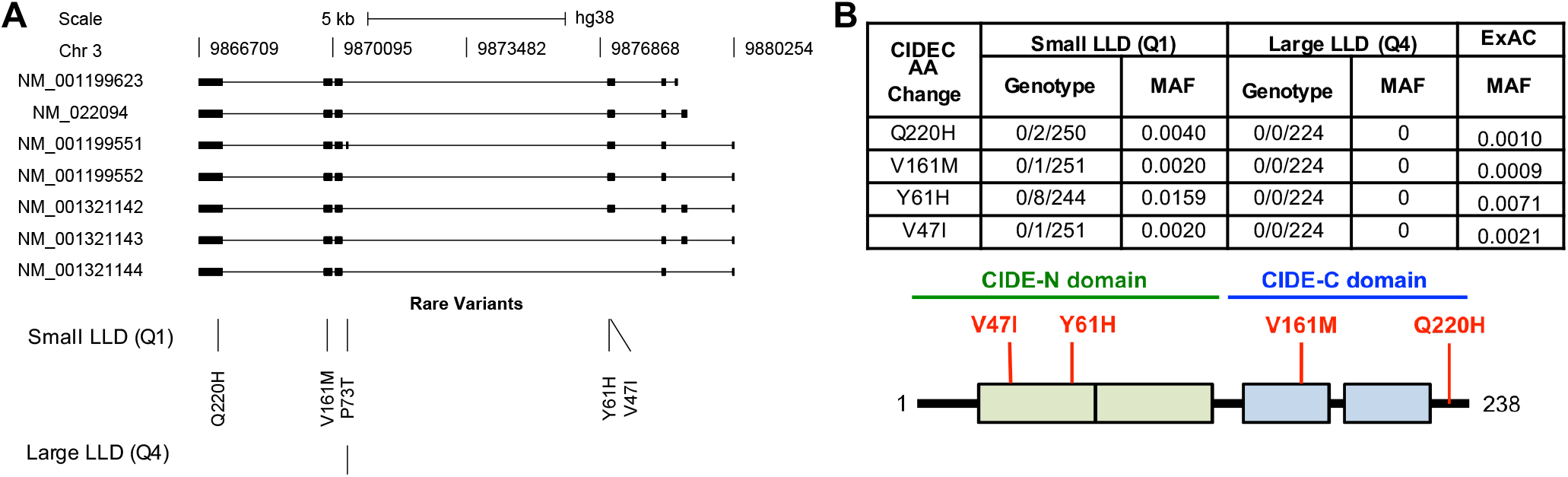
Genetic analysis of low-luminance deficit quartile Q1 and quartile Q4 AMD patients. (A) Genetic diagram of *CIDEC* and location of rare variants in Q1 and Q4 AMD patients. SNPs are indicated by amino acid change and position. (B) Table of genotype and minor allele frequencies for variants selected for further analysis and map of CIDEC protein with CIDE-N and CIDE-C domains with these SNPs annotated by position and amino acid change.

### Q1 AMD CIDEC rare alleles cause a defect in lipid droplet fusion and enlargement

*CIDEC* is a homolog of the murine *Fsp27* (Fat-specific protein 27kDa) gene ^20^. *Fsp27* was originally identified as a gene up-regulated during murine pre-adipocytes differentiation *in vitro* ^21; 22^. FSP27 was then shown to localize to lipid droplets (LDs) in adipocytes, where it promotes triglyceride storage by inhibiting LD fragmentation and lipolysis ^23^. *In vivo*, FSP27 is mainly expressed in the white adipocytes where it contributes to optimal energy storage by allowing the formation of their characteristic large unilocular LD ^24^. *Fsp27* deficient mice have white adipocytes with small multilocular LDs and increased mitochondrial size and activity, resulting in smaller white fat pads and increased metabolic rate ^24; 25^. A *CIDEC* homozygous nonsense mutation was identified in a patient with partial lipodystrophy and insulin resistant diabetes (OMIM: 615238) ^26^. This p.Glu186* (E186X, c.556G → T) mutation results in truncation of the CIDEC protein and the patient presented with multilocular small LDs and focal increased mitochondria density in adipocytes. Notably, *Fsp27* deficient mice have a healthy metabolic profile but when challenged by substantial energetic stress, they acquire features found in the CIDEC E186X patient, such as insulin resistance and hepatic steatosis ^27^. However, no eye phenotype has been reported in the CIDEC E186X patient nor the *Fsp27* deficient mice. Therefore, we first investigated the potential functional consequences of the four rare, protein altering CIDEC alleles found in Q1 AMD patients in adipocytes, a cell type in which CIDEC’s function has been well established.

First, we transiently expressed different versions of CIDEC tagged with GFP into 3T3-L1 pre-adipocytes. We transfected each of the four Q1 AMD rare variants (V47I, Y61H, V161M and Q220H) and as controls, we transfected cells with CIDEC wild-type (WT) or with the CIDEC E168X mutation. Subsequently, the proteins encoded by the Q1AMD rare *CIDEC* alleles will be referred to as “AMD CIDEC variants”. The cells were then treated for two days with oleic acid to induce LD formation. As expected, the mutant CIDEC E168X was diffused in the cytoplasm and failed to accumulate around the LDs (data not shown, and ^26^). In contrast, the four AMD CIDEC variants mostly localized to LDs in transfected adipocytes, and similarly to CIDEC WT, accumulated at the LD-LD contact sites (**Figure 3A**).

**Figure 3.**
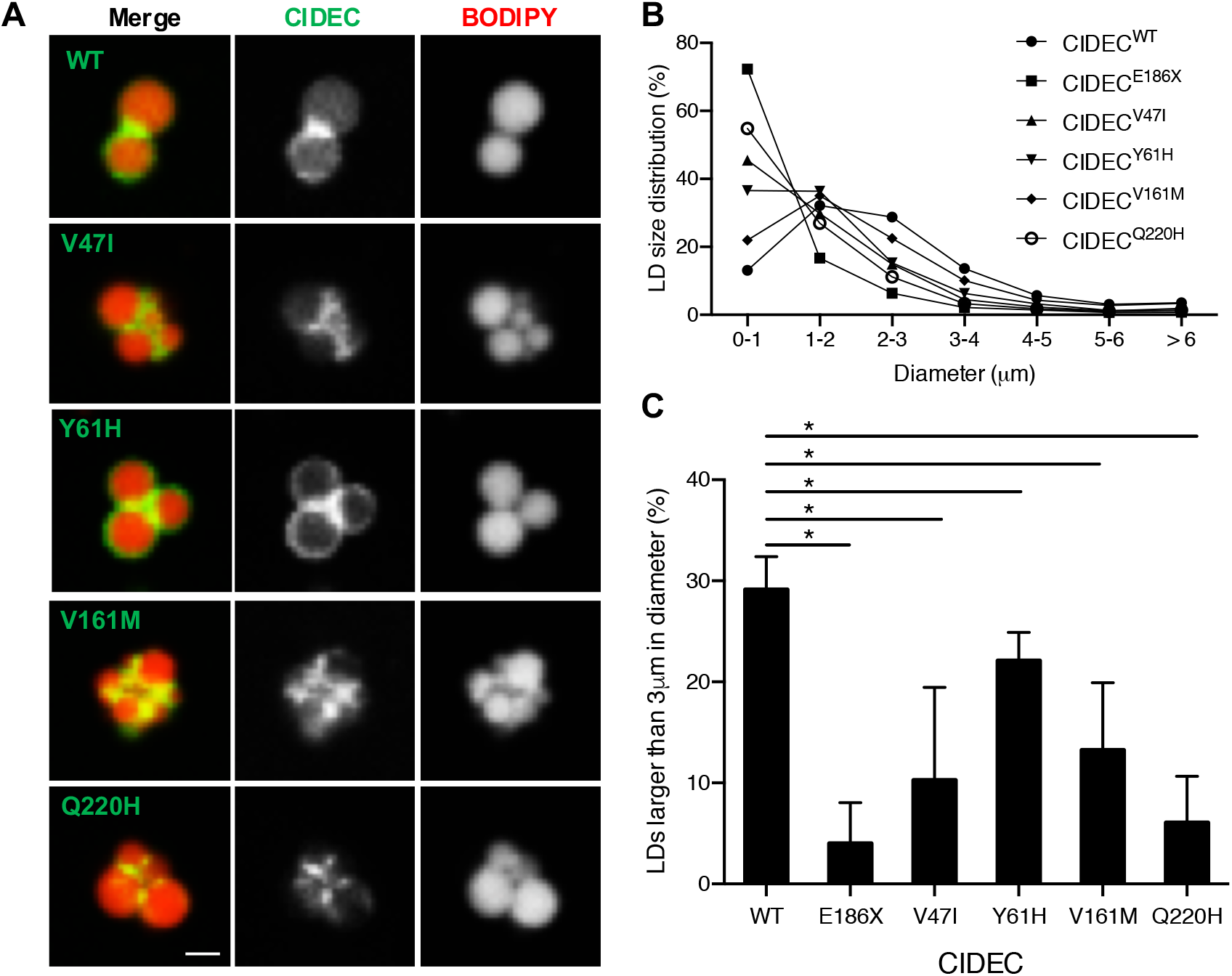
AMD CIDEC variants localize to lipid droplets (LDs) but cause a defect in LD enlargement. (A) Representative images of GFP-tagged CIDEC wild-type (WT) or rare variants localized to LDs labeled in red by BODIPY 558/568. Scale bar: 2 μm. (B) Size distribution of LDs in pre-adipocytes expressing CIDEC WT or each of the rare variants (diameters in μm). (C) Percentage of LDs with a diameter larger than 3 μm. N=3 (mean ± SD, Student’s t test, *p<0.05)

Next, we assessed the size of the LDs in the transfected adipocytes. We found that cells transfected with CIDEC WT had LDs with an average diameter of 2 to 3 μm. However, cells expressing CIDEC E168X had a severe LD enlargement defect, resulting in accumulation of clustered LDs with diameters smaller than 1 μm. Cells expressing each of the AMD CIDEC variants had an intermediate phenotype with a majority of LDs being smaller than 2 μm (**Figures 3B and C**). Interestingly, unlike in the E168X mutation case or *Fsp27* deficiency, we found that the presence of the AMD CIDEC variants did not increase the density of mitochondria in the transfected cells (**Figure S3A**) and they did not alter mitochondria activity as measured with a Seahorse bioanalyzer (**Figure S3B**). In conclusion, the AMD CIDEC variants do not impair proper CIDEC localization to LDs and do not increase mitochondrial density, but they are hypomorphic variants reducing the LD enlargement capacity in adipocytes.

Next, we transiently co-expressed GFP-tagged version of CIDEC WT or each of the four AMD CIDEC variants with mCherry-tagged Perilipin1 (PLIN1) in 3T3-L1 pre-adipocytes. PLIN1 is an adipocyte-specific LD-associated protein that interacts with and potentiates the function of murine CIDEC and hence could be used to track individual LDs ^28^. After inducing LD formation with oleic acid treatment, we performed time-lapse imaging over 6 hours to quantify the number of LD fusion events (**Figure 4 and Video 1**). Cells expressing each of the AMD CIDEC variants showed significant defects in LD fusion frequency compared to cells expressing CIDEC WT (Student’s t-test, p<0.005). Over 6 hours, cells expressing CIDEC WT had 14.6%±1.9% of their LDs achieving fusion (**Figures 4A and B**). Cells expressing CIDEC V47I, Y61H and V161M had a severe decrease in LD fusion events with only 0.3%±0.5%, 2.0%±1.8% and 1.2%±1.1% of LDs achieving fusion respectively. No LD fusion events were recorded during the 6 hours in cells expressing CIDEC Q220H, suggesting that this variant causes a severe loss of LD fusion capacity. Quantification of the time required from initial contact to complete fusion of two LDs revealed that LD fusion events slowdown in presence of the CIDEC variants (**Figure 4C**). In conclusion, adipocytes expressing the AMD CIDEC variants have a defect in LD fusion capacity, with merging events being slower and rarer than the ones occurring in cells expressing CIDEC WT.

**Figure 4.**
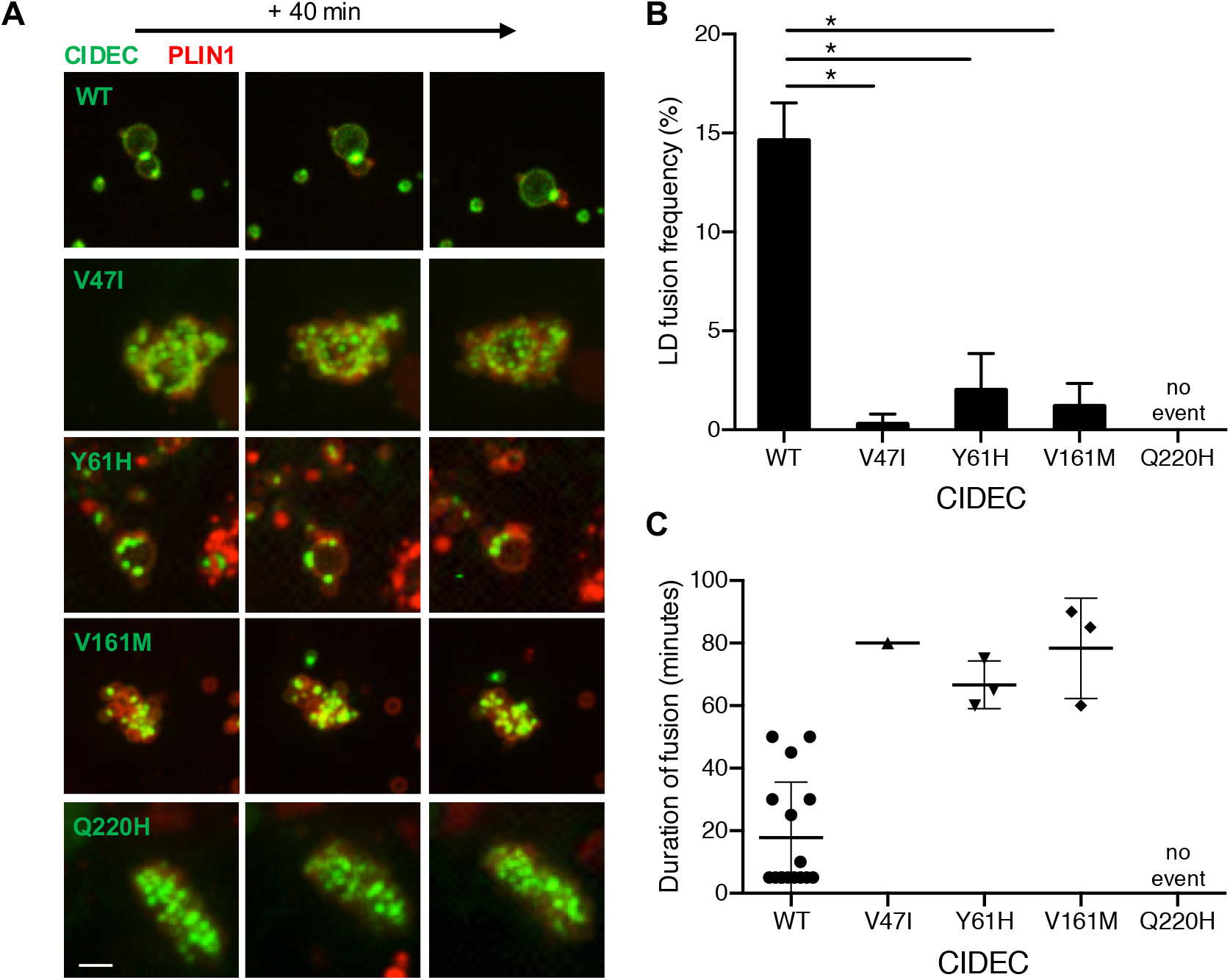
Lipid droplet (LD) fusion occurs less frequently and more slowly in pre-adipocytes expressing the AMD CIDEC rare variants. (A) Representative time-lapse images over 40 minutes of LDs in cells co-expressing GFP-tagged CIDEC WT (taken from **Video 1**) or each of the rare variants, and mCherry-tagged PLIN1. Scale bar: 2 μm. (B) Percentage of LDs undergoing fusion during the 6-hour analysis. N=3 (mean ± SD, Student’s t test, *p<0.05). (C) Time in minutes required from initial LD–LD contact to complete LD fusion.

Finally, we performed a Fluorescence Recovery After Photobleaching (FRAP) experiment to determine if the CIDEC variants could affect the kinetics of lipid diffusion between LDs. We transiently transfected 3T3-L1 pre-adipocytes with GFP-tagged CIDEC WT or AMD variants, induced LD formation and labeled LDs with the fluorescent fatty acid BODIPY 558/568 dye. Focusing on adjoining LDs of equivalent size and expressing CIDEC at the contact site, we photobleached one LD and measured the mean optical intensity (MOI) of both the bleached and the neighboring, unbleached LD. Recovery of fluorescence on the bleached LD over time was used to quantify the rate of lipid exchange between the two LDs (**Figure 5A**). In cells expressing CIDEC WT, the fluorescence recovered to about 75% of the pre-bleach intensity within 3 minutes in the photobleached LD. This recovery was accompanied by a corresponding decrease in fluorescence on the unbleached LD, reflecting efficient lipid exchange between the two LDs (**Figures 5B and C**). In cells expressing CIDEC Y61H or V161M, fluorescence recovery in the bleached LD exhibited a delayed and reduced fluorescence compared to cells expressing CIDEC WT. In cells expressing CIDEC V47I or Q220H there was very limited, if any, fluorescence recovery on the bleached LD, suggesting loss of lipid exchange capacity (**Figures 5B and C**). In conclusion, the presence of the AMD CIDEC variants impairs the lipid diffusion capacity between LDs in adipocytes.

**Figure 5.**
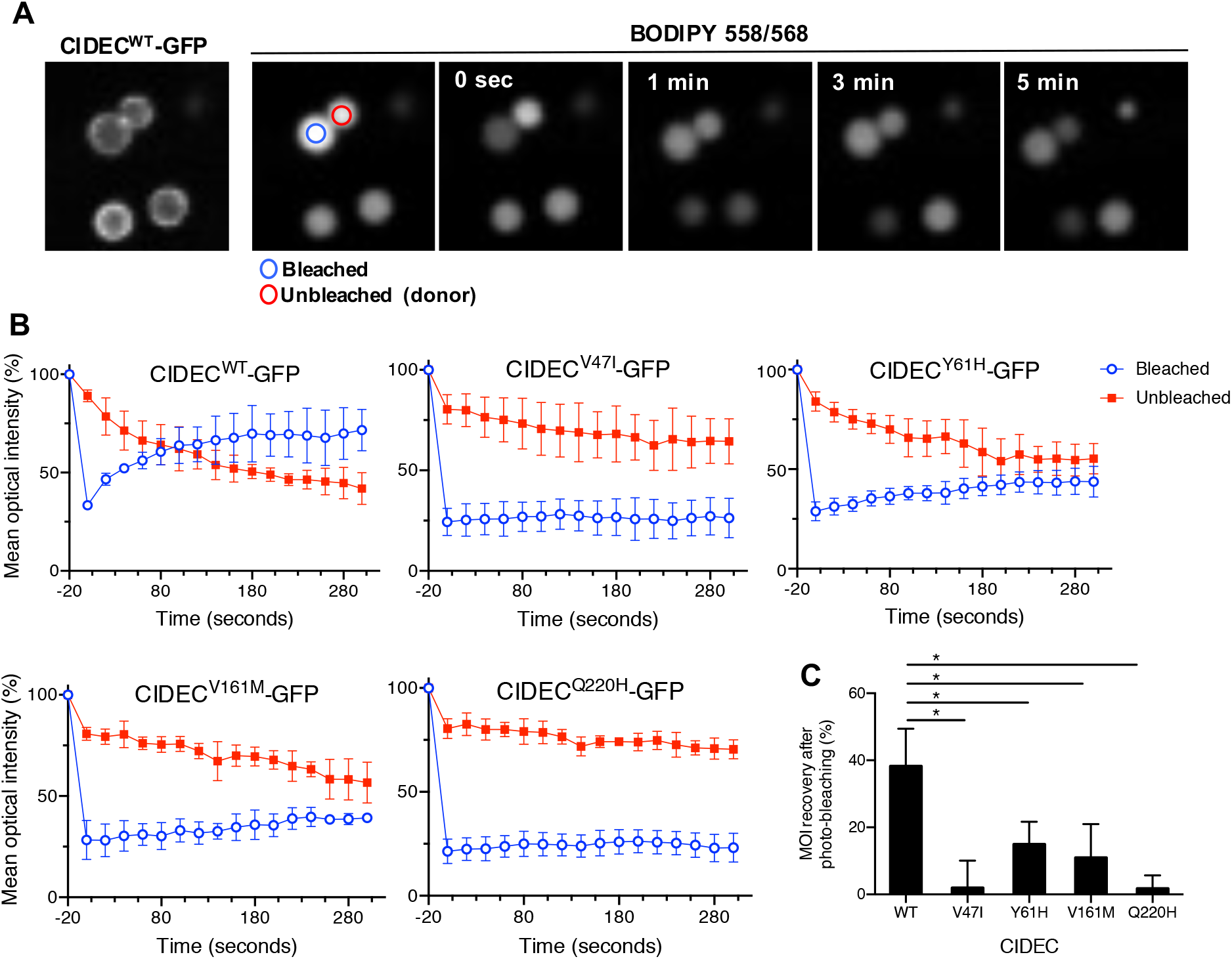
AMD CIDEC variants cause a decrease in the lipid exchange rate between lipid droplets (LDs). (A) Representative Fluorescence Recovery After Photobleach (FRAP) images of paired LD expressing GFP-tagged CIDEC wild-type (WT) showing progressive neutral lipid (BODIPY 558/568 dye labeling) exchange as determined by fluorescence recovery from the adjacent LD. (B) Quantification of mean optical intensity (MOI) in the bleached (blue circle) and unbleached (red circle) LD in cells expressing CIDEC WT or each of the rare variants. (C) Percentage of MOI recovery on bleached LDs from 0 sec. to 300 seconds. N=3 (mean ± SD, Student’s t test, *p<0.05).

Collectively, these results show that the AMD CIDEC variants do not affect CIDEC localization to LD contact sites, but they impair the lipid exchange capacity between LDs, resulting in defective LD fusion and incapacity for the adipocytes to accumulate lipids inside few large LDs.

### Q1 AMD CIDEC rare alleles decrease the binding affinity of CIDEC with the lipid droplet fusion effectors PLIN1, RAB8A and AS160

LDs are highly dynamic organelles containing a neutral lipid core enclosed in a phospholipid monolayer decorated by a large number of proteins ^29; 30^. To better understand the functional consequences of the AMD CIDEC alleles and how they can affect LD fusion, we examined if they could alter protein-protein interactions. Indeed, CIDEC-mediated LD fusion is different from other membrane fusion events. CIDEC proteins need to first accumulate at the contact site between two LDs, to then enable recruitment of regulator proteins such as PLIN1, RAB8A and AS160, which facilitate the lipid transfer through the fusion pore ^31^.

We first assessed if the variants affected CIDEC capacity to homodimerize. CIDEC contains two conserved CIDE domains allowing its dimerization, a N-terminal CIDE-N domain and a C-terminal CIDE-C domain (^20; 32^ and **Figure 2B**). The CIDE-N domain, in which the variants V47I and Y61H are located, dimerizes mainly via electrostatic interactions, while the CIDE-C domain that contains the V161M variant dimerizes through a stronger interaction ^28^. Q1 AMD patients are heterozygous for the different CIDEC alleles, so HEK 293T cells were co-transfected with 3xFlag-CIDEC WT and either CIDEC WT, the E186X mutation or one of the AMD CIDEC variants tagged with GFP. After immunoprecipitation of the 3xFlag-CIDEC WT, pulled-down proteins were probed with anti-GFP. Co-transfection of the 3xFlag-CIDEC WT together with GFP alone served as negative control. The CIDE-N domain variants V47I and Y61H showed decreased dimerization capacity with CIDEC WT, whereas the two other variants V161M and Q220H did not affect the binding ability (**Figure 6A**). The pathogenic mutation CIDEC E186X, located in the CIDE-C domain, also did not affect the binding affinity with CIDEC WT.

**Figure 6.**
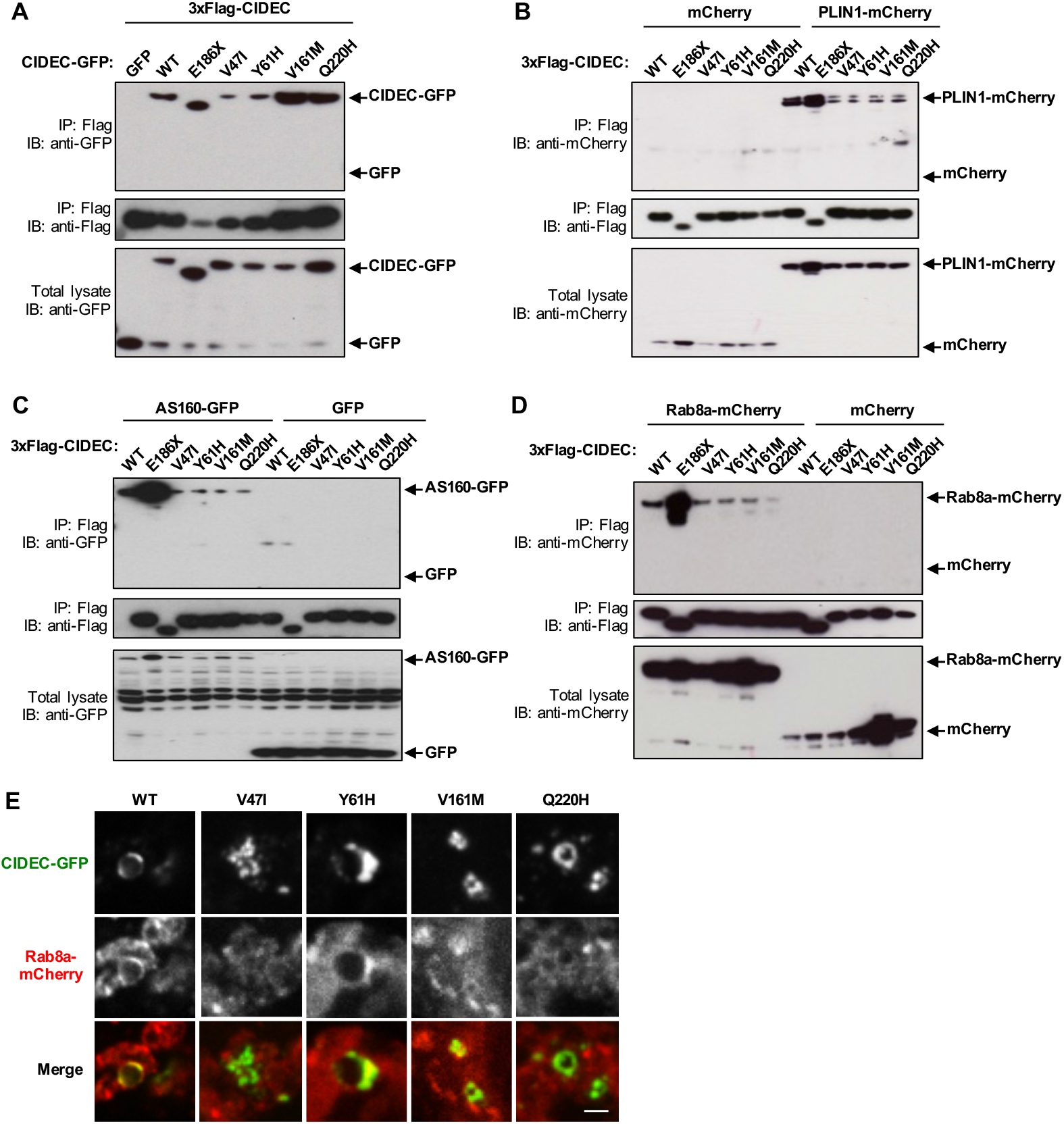
AMD CIDEC variants in the CIDE-N domain decrease dimerization affinity and all four variants decrease binding to effector partners PLIN1, AS160 and RAB8A. (A) 3xflag-tagged CIDEC wild-type (WT) was co-transfected with the indicated GFP-tagged CIDEC variants in HEK 293T cells. GFP alone was used as negative control. 3xflag-tagged CIDEC WT was immuno-precipitated (IP) using anti-Flag and pulled-down proteins were immuno-blotted (IB) with anti-GFP and anti-Flag. Total cell lysate was immunoblotted with anti-GFP to control for CIDEC-GFP expression levels. (B-E) HEK 293T cells were co-transfected with 3xFlag-CIDEC WT, E186X or AMD variants, and either PLIN1-mCherry (B), AS160-GFP (C) or RAB8A-mCherry (D). After immunoprecipitation (IP) of the 3xFlag-CIDEC, pulled-down proteins were probed with anti-mCherry or anti-GFP, and anti-Flag. Co-transfection with mCherry or GFP alone was used as negative controls. Total cell lysates were immunoblotted (IB) with anti-mCherry or anti-GFP to control for PLIN1, AS160 and RAB8A expression levels. (E) Representative fluorescence images of 3T3-L1 pre-adipocytes lipid droplets containing CIDEC-GFP wild-type (WT) or variants and RAB8A-mCherry. Scale bar: 2 μm.

Next, we assessed if the AMD CIDEC variants affect CIDEC capacity to interact with its LD-associated regulatory partners PLIN1, RAB8A and AS160, as these interactions are required for LD fusion and growth ^28; 33^. We co-transfected HEK 293T cells with 3xFlag-CIDEC WT, E186X or the AMD CIDEC variants, and either PLIN1-mCherry, AS160-GFP or RAB8A-mCherry. After immunoprecipitation of the 3xFlag-CIDEC, pulled-down proteins were probed with anti-mCherry or anti-GFP. Strikingly, all four AMD CIDEC variants had similarly decreased binding affinity with PLIN1 (**Figure 6B**) and AS160 (**Figure 6C**). All four AMD CIDEC variants also showed decreased binding capacity with RAB8A, however, the Q220H variant caused a more severe loss of interaction with the GTPase (**Figure 6D**). Only a fraction of RAB8A and AS160 are associated to LDs, with the rest being distributed in the cytoplasm (**Figure 6E and Figure S4**). The fact that the E186X mutant is abnormally diffuse in the cytoplasm could explain its stronger interaction with the binding partners compared to CIDEC WT, which is concentrated on the LDs (**Figures 6B, C and D**).

Collectively, these results show that the AMD CIDEC variants V47I and Y61H, located in the CIDE-N domain decrease CIDEC dimerization capacity and its binding ability with the regulators partners PLIN1, RAB8A and AS160. The two other variants, V161M and Q220H, which are not in the CIDE-N domain, do not affect CIDEC ability to dimerize, however, they nevertheless also decrease its interaction with PLIN1, RAB8A and AS160. The reduced interaction capacity of the four AMD CIDEC variants with its binding partners may explain how their presence causes a defect in lipid droplet fusion and enlargement in adipocytes.

### CIDEC expression is not detected in the human retina or the Retinal Pigment Epithelium and the Q1 AMD CIDEC variants do not affect the size of the retinosomes

CIDEC plays a critical role in the white adipose tissue, but is also expressed in organs such as muscles, nerves and even blood vessels ^18^. CIDEC expression in the eye has been reported after an Expressed Sequence Tag (EST) database search, however, it is not known if the expression comes from the neuroretina or the eye globe supportive tissue ^32^. The key elements of the eye involved in AMD are the photoreceptors, an epithelium located underneath the retina called the Retinal Pigment Epithelium (RPE) and the blood vessels supporting the retina called the choroid. To investigate the potential expression of *CIDEC* in these structures, we first used published human RNA sequencing datasets. *CIDEC* was not detect in human retina or RPE/choroid (bulk RNA sequencing^34^) and in different human ocular cell types (single cell RNA sequencing^35^) (**Figure 1C**). To investigate further the potential expression of *CIDEC* in the eye, we performed RNA *in situ* hybridization on eye sections from a Caucasian 73-year old female and a Caucasian 88-year old male, both without history of AMD (**Figure 7A**). We also performed *Fsp27* RNA *in situ* hybridization on mouse eye sections (**Figure S5**). In both human and mouse eyes, we did not detect CIDEC RNA in the retina, the RPE or the choroid (**Figure 7A and Figure S5**). We used the sensitive detection method BaseScope™ (Advanced Cell Diagnostics (ACD)) and found that the signal detected with the CIDEC probes on the human eye sections were consistent with the background signal detected using the bacterial gene DapB as a negative control (**Figure 7A**). On the mouse eye sections, we found rare cells positive for Fsp27 expression but these cells were in the supportive tissue around the eye (**Figure S5**).

**Figure 7.**
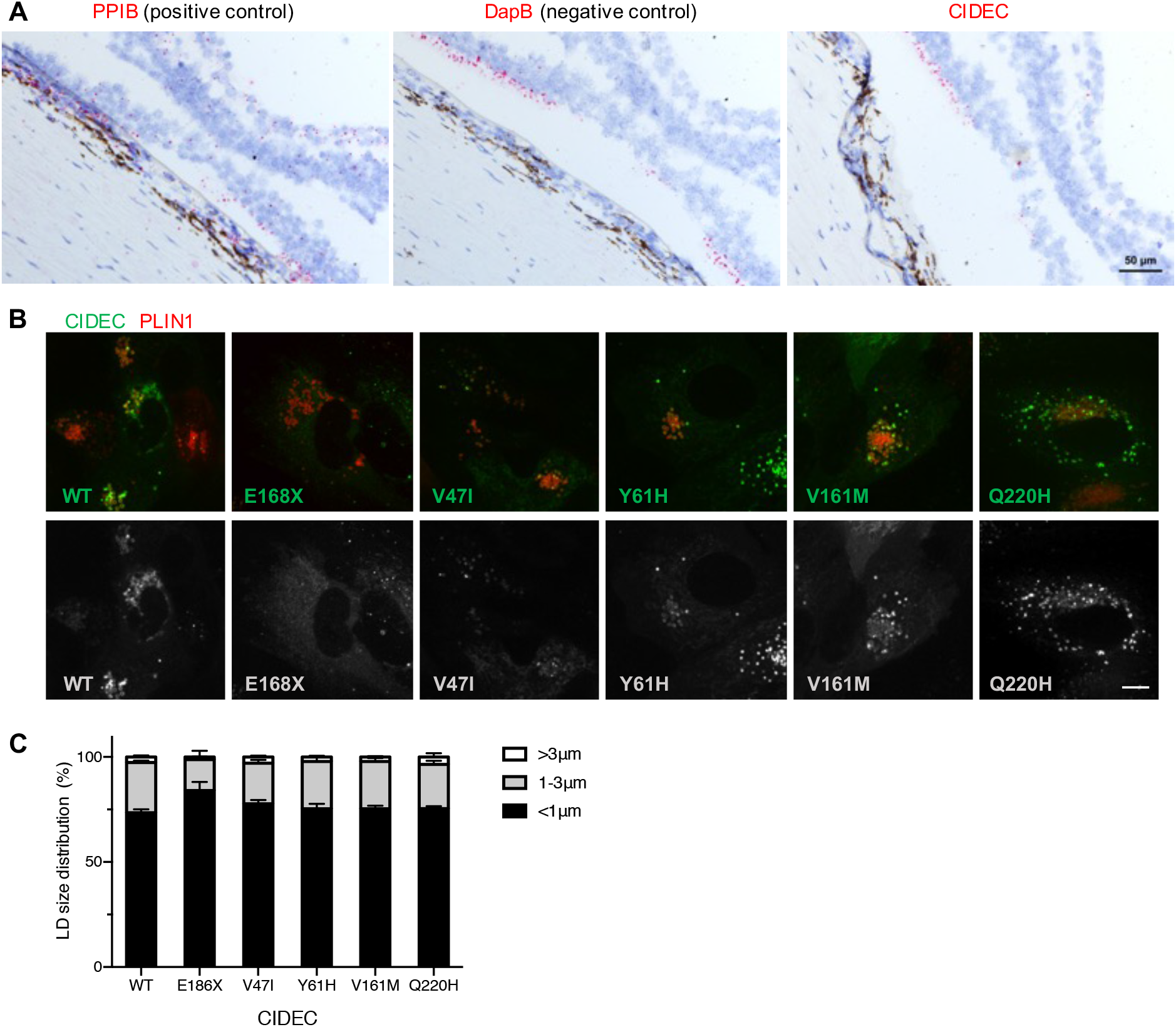
CIDEC RNA is not detected in the human eye and exogenous expression of the CIDEC variants does not affect lipid droplets (LDs) size in Retinal Pigment Epithelium (RPE) cells. (A) In situ hybridization in the fovea of a control human donor eye showing that CIDEC RNA is not detected in the retina or RPE cells. Detection of PPIB (red) was used as positive control and detection of bacterial DapB was used as negative control and evaluation of the non-specific background. Scale bar: 50 μm. (B and C) Human fetal RPE cells were co-infected with lentivirus expressing CIDEC variants and PLIN1 as marker for LDs. The infected cells were differentiated for 3 weeks before oleic acid stimulation. Representative images of RPE cells expressing both CIDEC variants and PLIN1 (B). LD diameters were quantified by diameter range as depicted in the bar graph (n=3; mean ± SD) (C). Scale bar: 5 μm.

RPE cells contain specific LDs called retinosomes, in which retinyl esters are stored and used to replenish key components of the visual cycle ^36; 37^. To account for the possibility that CIDEC was expressed below our detection threshold in RPE cells, we tested if exogenous AMD CIDEC variants could have consequences on the size of these specialized LDs, retinosomes. Primary human fetal RPE cells were infected with lentivirus encoding CIDEC WT or the AMD CIDEC variants and differentiated for three weeks before oleic acid stimulation and LD labelling. Similar to the localization in adipocytes, CIDEC WT and AMD CIDEC variants accumulate on retinosomes and concentrate at the LD fusion sites in RPE cells (**Figure 7B**). However, the RPE cells failed to form large LDs after oleic acid stimulation and the majority of the retinosomes in RPE cells expressing CIDEC WT were smaller than 1 µm in diameter (**Figure 7C**). Consequently, we did not observe any difference in LD size between the RPE cells expressing the CIDEC WT and the cells expressing the different AMD CIDEC variants.

Finally, we compared color fundus photos (**Figure 8**) and Optical Coherence Tomography (OCT) images (not shown) from the eyes of the Q1 AMD CIDEC variant carriers and Q1 AMD CIDEC variant non-carriers. In particular, we wanted to know if by disrupting lipid accumulation, the CIDEC variants could affect size and accumulation of drusen, which are deposits of proteins and lipids building up under the retina and a hallmark of AMD. However, we did not observe unique ocular clinical features in patients carrying the CIDEC rare variants (**Figure 8B**) compared to non-carriers (**Figure 8A**). In both groups, we observed typical AMD clinical features such as pigmentary changes, variable amount of drusen, geographic atrophy and choroidal neovascular lesions.

**Figure 8.**
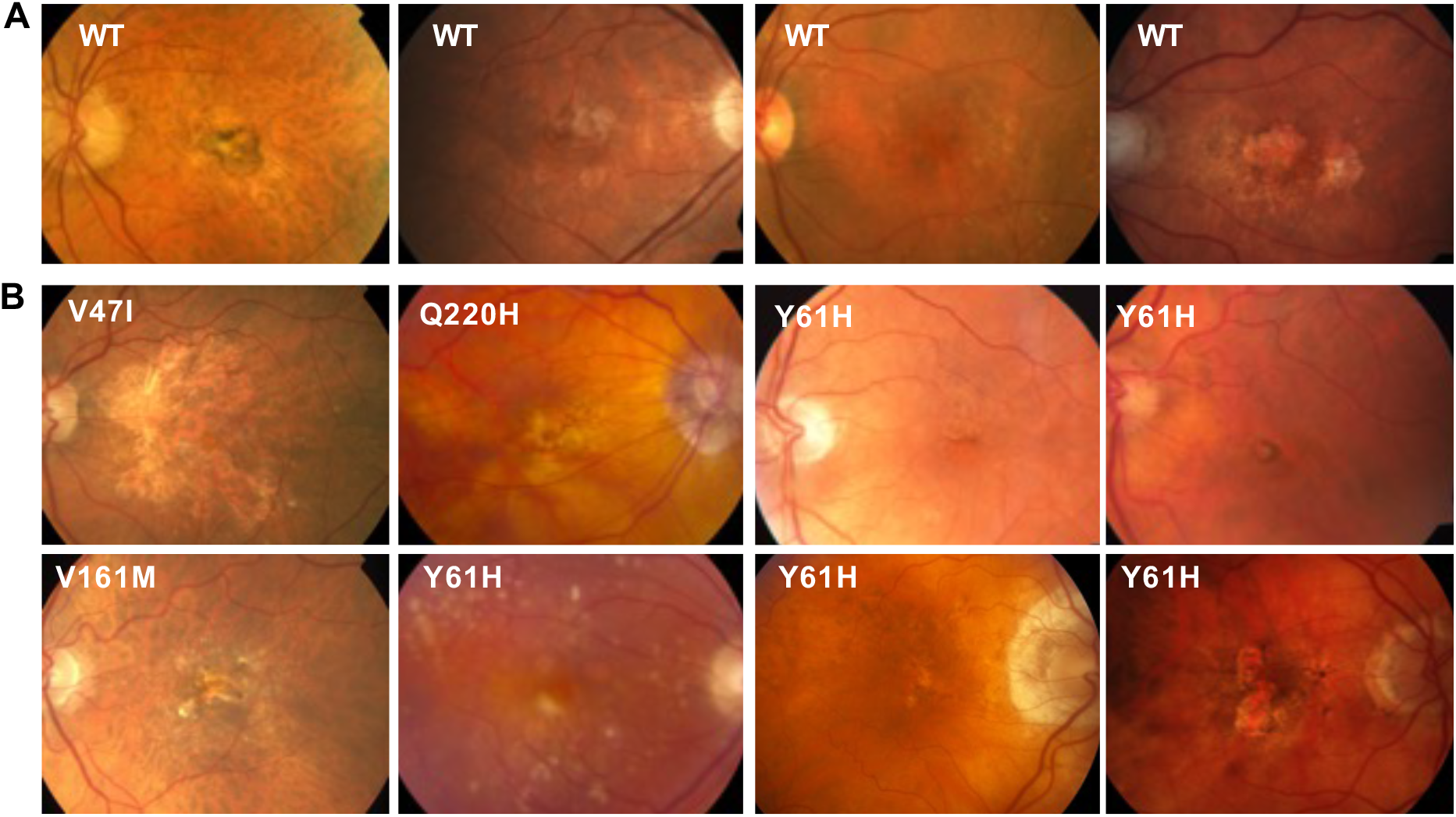
Clinical images of patients in the low-luminance deficit quartile Q1. Color fundus photos (CFP) from participants in the HARBOR trial (A) Q1 non-carriers for CIDEC rare variants (B) Q1 rare variant CIDEC carriers. CFP in both groups demonstrate typical clinical features of macular degeneration such as pigmentary changes, drusen, geographic atrophy and choroidal neovascular lesions. No obvious phenotypic differences are noted between the two groups.

In conclusion, we did not detect CIDEC expression in the ocular structures directly affected in AMD. We also found that exogenous expression of the AMD CIDEC variants did not alter retinosome size in RPE cells, and that AMD patients carrying the CIDEC variants do not present unique phenotypic ocular features compared to non-carriers. Our results suggest that the AMD CIDEC variants do not play a direct role in the eye. Additional experiments using conditional mouse models will be important to assigning the tissue specific effects of CIDEC variants and the role of LD dysregulation in AMD.

## Discussion

Here we report the first analysis examining the genetic effect on baseline LLD, a clinical measurement that has been shown to be predictive of anti-VEGF treatment response and GA lesion growth, in AMD patients. While the study is of modest size, to our knowledge, it is novel in its effort to utilize clinical indices beyond BCVA that have been linked to patient outcomes to further homogenize AMD patients in order to increase power for genetic analysis. It is our hope that as datasets increase in size and have deeper phenotypic assessment, these types of sub-phenotype GWAS analyses will increase and work alongside recent studies utilizing novel *in vitro* methods, such as those described here and genome-wide single cell and perturbation methods to help uncover the functionality of genes associated with the pathogenesis of AMD.

We did not find any variants, either in the common or the rare variant burden analysis which passed a pre-specified significance threshold accounting for multiple testing. This can occur for several reasons. One reason could be that the effect of patient germline genetics is not substantial on low-luminance visual acuity and that environmental factors explain more of the risk variability. Another is that we are underpowered for a genome wide analysis in our study population. As such, it will be important to replicate the genetic findings in a secondary cohort which has similar phenotyping and sequencing data. Since all patients in our analysis are advanced AMD patients, and *CIDEC* has not been reported previously as an AMD risk gene, this could indicate that *CIDEC* rare variants play a role only once a patient develops advanced disease. Datasets with deep phenotyping of advanced AMD patients would be required for replication. While large scale biobank data exist (e.g. U.K. BioBank), and are exceptionally useful for most replication analyses, these datasets do not currently have the ability to delve deep into clinical features for an age-related disease such as AMD. Conversely, smaller, more phenotypically focused genetic datasets such as the one used in this study are useful for identification of signals and hypotheses, but are severely underpowered to confirm an association statistically. As such, we sought to assess the possible contribution of *CIDEC*, a gene with biology tangential to genes in known AMD risk loci, to AMD pathology through *in vitro* analysis.

In our rare variant burden analysis, we noticed that one of the top hits was strongly linked with lipid metabolism. Similarly, multiple loci identified in AMD risk studies contain genes implicating lipid metabolism. Due to sample size constraints, and the low frequency of CIDEC variants, we were unable to test for interaction between known AMD risk alleles and the CIDEC variants. Rare, protein altering variants were enriched in the patients with small LLD. Previous studies associated a small LLD at baseline with more favorable prognostic and predictive outcomes. Thus, possible impairment of CIDEC function could be beneficial for AMD patients and we undertook characterization of the *CIDEC* variants in further studies, which focused on the exact variants seen in our AMD cases.

In our *in vitro* analysis, we interestingly found that all four rare variants failed to impair CIDEC localization to LDs, but instead all decreased the binding affinity of CIDEC with the LD fusion effectors PLIN1, RAB8A and AS160. Interaction of CIDEC with these binding partners is critical for its function and we hypothesize that this decreased interaction underlies the defect in LD enlargement and lipid exchange that we observed in adipocytes expressing the AMD CIDEC variants. Interestingly, the functional consequences that we uncovered are milder than the ones caused by the lipodystrophic E168X variant and are restricted to the LD capacity to fuse and increase lipid storage. Indeed, the AMD CIDEC variants are hypomorphic regarding LD size and they do not affect mitochondria density or activity. Our data suggest that the Q1 AMD CIDEC variants do not severely disrupt adipocyte health and function and may have a beneficial effect by only limiting the capacity of adipocytes to accumulate lipids in very large LDs.

Of note, patients carrying the Q1 AMD CIDEC variants are heterozygotes, contrasting with the E168X homozygote lipodystropic patient. Furthermore, heterozygous Fsp27 wt/ko mice have normal weight and the appearance of their adipose tissue is similar to the one from Fsp27 wt/wt mice ^24^. Thus, it is likely that the Q1 AMD CIDEC patients did not suffer from severe lipodystrophy and that they only had sub-clinical consequences of the CIDEC variants expression. Our results suggest that CIDEC is not expressed in the ocular tissue affected in AMD such as the retina, RPE and choroid. This points out toward a “systemic” effect of the beneficial Q1 AMD CIDEC variants. Interestingly, a similar indirect and systemic favorable effect has recently been reported in mouse models of vascular inflammation and atherogenesis after *Fsp27* silencing^38; 39^. Many studies on dietary or circulating lipids, as well as genetic studies support a role for not only local lipid trafficking in the retina but also for circulating lipoproteins in AMD pathogenesis ^40^. Therefore, it would have been interesting to perform a biomarker investigation of the HARBOR patient serum to know if any particular change(s) in circulating lipoproteins levels could be detected in patients carrying the Q1 AMD CIDEC variants compared to non-carriers (unfortunately, such samples were not available for us to perform the analysis). In addition to lipoproteins, it would also have been interesting to probe the AMD patient serums for changes in adipokines, the cytokines secreted by adipocytes. Indeed, it has been shown that hypertrophied adipocytes can lead to local inflammation and inhibition of production of adipokines, such as adiponectin ^41^. Since adipocytes expressing the AMD CIDEC variants have a decreased LD enlargement capacity, it may prevent them from becoming hypertrophic, keeping adiponectin level high. Supporting this idea, it has been reported that the *Fsp27* deficient mice show increased serum adiponectin level compared to wild type mice ^27; 42^. Importantly, increased serum adiponectin levels have been shown to be protective in several pre-clinical models of angiogenesis in the eye, including models of neovascular AMD ^43–45^ and there is human genetic data linking adiponectin ADIPOQ and its receptor ADIPOR1 to the risk of advanced AMD ^46; 47^. Finally, CIDEC, ADIPOQ and APOE (an AMD GWAS locus also involved in lipid metabolism) have been linked as part of an 8-gene hub identified as candidate serum biomarkers for diabetic peripheral neuropathy^16^.

In conclusion, our rare variant burden genetic analysis followed by our *in vitro* dissection of the functional consequences of the beneficial variants, altogether with published data, suggest that once patients have developed advanced AMD, the disease outcome could be modified by systemic and indirect lipidomic biological processes that it would be interesting to investigate further. In particular, investigating adipokines serum level, including adiponectin, in advanced AMD patients could provide new biomarkers of neovascular AMD progression or response to anti-VEGF therapy.

## Materials and Methods

### Research subjects and low-luminance deficit (LLD)

Before execution of the study, an internal Genentech team of Informed Consent Form (ICF) experts reviewed the ICFs from all the studies to ensure appropriate use of the samples. The HARBOR clinical trial (ClinicalTrials.gov identifier: NCT00891735) was a 24-month Phase III study designed to evaluate the effectiveness of monthly or as-needed ranibizumab delivery in patients with subfoveal neovascular AMD. This study has been described previously^13; 14^. LLD dysfunction is quantified by first assessing best corrected visual acuity (BCVA) under normal lighting conditions, followed immediately by a low-luminance visual acuity (LLVA) measurement, and this has been described previously ^11; 12^.

### Patient population and genetic analysis

As we are looking at baseline characteristics, all HARBOR patients, regardless of randomized treatment assignment, were eligible for inclusion in our study. We excluded patients who did not consent for exploratory analyses and patients of non-European descent. This resulted in the removal of 118 individuals from the overall enrolled trial study population, representing 10.7% of the study population. We stratified patients for analysis based on LLD quartile 1 (Q1) vs quartile 4 (Q4) as described previously ^12^. This resulted in 275 patients in our Q1 group, and 241 patients in Q4. Further demographic information is found in **Table 1**. Logistic regression was used to assess the association in the common variant analysis, adjusted for age, sex, baseline visual acuity and genetically determined ancestry. PLINK version 1.90b3.46 was used for the common variant analysis.

The sequence data was annotated using SnpEff and there were 120,580 exonic coding variants at a minor allele frequency < 1%. For a gene to be included in the analysis, it had to contain at least two coding SNPs, resulting in 13,046 genes that could be tested for rare-variant gene-burden. The rare variant gene burden test was used to assess the cumulative effect of rare variants. Rvtest software (version 20170228) was used for the combined multivariate and collapsing gene burden test, adjusted for age, sex, baseline visual acuity and genetically determined ancestry. The rare variant gene burden test was used to assess the cumulative effect of rare variants (MAF < 1%).

### Cell culture and treatments

293T cells (ATCC #CRL-3216) and 3T3-L1 preadipocytes (ATCC #CL-173) were cultured in Dulbecco’s modified Eagle’s Medium (DMEM) containing 10% fetal bovine serum (FBS) and Penicillin (10,000 units/ml)/ Streptomycin (10,000μg/ml, 1:100 dilution of stock, Gibco #15140-122). After reaching confluency, 3T3-L1 pre-adipocytes were cultured for 48 hours in DMEM + 10% FBS. The culture medium was then replaced by DMEM + 10% FBS + 5 μg/ml insulin (Sigma, I0516) + 1 μM dexamethasone (G-bioscience, API-04) + 0.5 mM isobutylmethylxanthine (Sigma, I5879)] to induce adipocyte differentiation. After 48 hours, the medium was replaced by DMEM + 10% FBS + 5 μg/ml insulin for an additional 48-72 hours to achieve complete differentiation. To induce lipid droplet formation, cells were treated with 200 μM of oleic acid-albumin from bovine serum (Sigma, O3008). Human fetal retinal pigment epithelial cells (hfRPE, Lonza #00194987) were cultured in RtEGM with supplement medium as indicated by the manufacturer’s protocol (RtEGM bullet kit, Lonza, #00195409). HfRPE cells were cultured to high confluence on coverglass culture plates (Thermo, 155411) for three weeks to obtain polarized RPE monolayers. After differentiation, hfRPE cells were treated with 20μM A2E (*N*-Retinylidene-*N*-Retinylethanolamine, 20mM stock dissolved in DMSO, Gene and Cell Technologies) for 24 hours.

### Plasmids, transfection and viruses

3x Flag- and GFP-tagged expression plasmids were used to express human CIDEC wild type (WT) and the CIDEC rare variants (E186X, V47I, Y61H, V161M, or Q220H) (Genecopoeia, Inc.). GFP-and mCherry-tagged plasmids were used to express human PLIN1, AS160, and RAB8A (Genecopoeia, Inc.). 293T and 3T3-L1 cells were transiently transfected using Lipofectamine2000 and Lipofectamine3000 (Invitrogen, 11668 and L3000). Expression of GFP-tagged CIDEC in hfRPE cells was carried out using lentivirus infection. Viral media were collected from 293T cells transiently transfected with viral vector (expression plasmid), delta8.9, and VSV-G in a molar ratio of 1:2.3:0.2 using Lipofectamine2000. HfRPE cells were infected (without polybrene) on the day when they were split onto 6 well culture apparatuses and kept in viral media for 4-5 days.

### Immunofluorescence, Immunoprecipitation and Immunoblotting

Anti-Flag (Sigma, F7425), anti-GFP (Abcam, ab6556), anti-mCherry (Abcam, ab167453) antibodies were obtained from commercial sources. Alexa 488-, Alexa-594-conjugated secondary antibodies were obtained from Invitrogen. HRP-labeled secondary antibodies were purchased from Cell Signaling Technologies. Cells were fixed with 4% Paraformaldehyde (EMS, 15710S) for 15 minutes and mounted using ProLong Gold anti-fade mounting medium with DAPI (Thermo Scientific, P36941). Images were obtained with a Nikon A1R confocal microscope or Yokogawa CSU-X spinning disk on a Nikon TiE microscope and a Photometrics Prime 95B. Image acquisition was performed using the NIS elements software 4.50 (Nikon). Co-immunoprecipitation was performed on 293T cells lysed in IP Lysis buffer (Pierce #87788) containing a proteasome inhibitor cocktail (Pierce, Thermo Scientific, 87788) two days after transient transfection. The cell lysates were incubated with anti-Flag M2 affinity beads (Sigma, F2426) overnight at 4°C. After pull-down of the agarose beads, the immunoprecipitates were washed three times with IP Lysis buffer and eluted in a 2x BOLT Lithium dodecyl sulfate sample buffer for Western blot analysis. The samples were electrophoresed on NuPage 4–12% Bis-Tris gels (Invitrogen #NP0303) in MES-SDS running buffer (Invitrogen, #NP0002) and transferred to PVDF membrane (Invitrogen, #IB24001) for immunoblotting.

### Lipid droplet (LD) assays

3T3-L1 preadipocytes were fixed, stained with the LD marker Bodipy 558/568 C12 fatty acid (Molecular Probes, D3835) and LD diameters were measured in 100 to 150 cells from three independent experiments using Imaris software (Bitplane) and Matlab image processing toolkit. For live cell imaging, 3T3-L1 preadipocytes were transiently co-transfected with GFP-tagged CIDEC and PLIN1-mCherry as LD markers, and incubated with 200 μM of oleic acid. Images were taken using the Nikon TiE spinning disk confocal microscope with an environmental chamber (Okolab) for 12 hours in 5-minute intervals. The frequency of LD fusion per cell and the time duration of LD fusion from three independent experiments were quantified and plotted using Microsoft Excel 2011 and Graphpad Prism version 8.0.1. Fluorescence Recovery After Photobleaching (FRAP)-based lipid diffusion assays were conducted on the Nikon A1R confocal microscope. FRAP was performed on 3T3-L1 preadipocytes transiently transfected with GFP-tagged CIDEC were incubated with 200 μM of oleic acid and stained with Bodipy 558/568 C12 fatty acid (Molecular Probes, D3835) for 15 hours. One hour before the beginning of the FRAP assay, the medium was changed. LD pairs with clear GFP expression at the contact sites were selected for bleaching. Selected regions were bleached with a 561 mm laser at 100% power for 62.4 milliseconds, followed by time-lapse scanning of 20-second intervals. Mean optical intensity (MOI) of the bleached and the unbleached adjacent LD was measured by ImageJ and plotted using Microsoft Excel 2011 and Graphpad Prism version 8.0.1.

### Mitochondria assays

3T3-L1 cells expressing GFP-tagged human CIDEC were incubated with MitoTracker (#M7512; Thermo Fisher Scientific) before fixation, followed by permeabilization with 0.5% Triton X-100 and staining with DAPI. The mitochondrial density of the CIDEC-expressing cells was determined by measuring the fluorescent intensity of the MitoTracker signal using ImageJ. For the Seahorse Cell Mito Stress Test, 3T3-L1 cells expressing GFP or GFP-tagged CIDEC WT and rare variants (V47I, Y61H, V161M, Q220H, and E186X) were plated on a 96-well assay plate (10^4^ cells/well). The cells were maintained in XF assay medium (Agilent, #102365100) and subjected to a mitochondrial stress test, using the extracellular flux assay kit by sequentially applying oligomycin (2 mmol/L), carbonyl cyanide 4-(trifluoromethoxy) phenylhydrazone (FCCP; 5 mmol/L), and antimycin/rotenone (1 mmol/L and 1 mmol/L) (Cell Mito Stress Test Kit, Agilent, #103015100). Analysis was carried out by using the Seahorse analyzer software.

### In situ hybridization

The *in situ* hybridization (ISH) BaseScope™ v2 assay (Advanced Cell Diagnostics (ACD)) was performed on 5 μm-thick formalin-fixed paraffin-embedded sections of adult human eyes according to the BaseScope™ detection reagent kit v2 ACD protocol. Probes against the ubiquitously expressed isomerase PPIB were used as positive control, and probes against bacterial DapB were used as negative control. Six custom probes of 18–25 bp oligonucleotide sequences were designed by ACD for highly specific and sensitive detection of human CIDEC RNA. After deparaffinization in xylene and endogenous peroxidase activity inhibition by H_2_O_2_ (10 min), sections were permeabilized and submitted to heat (15 min at 100°C) and protease IV treatment (20 min at 40°C). After probe hybridization for 2 hours at 40°C, the signal was chemically amplified using the kit reagents and detected using the FastRED dye. The sections were then counterstained with Hematoxylin and mounted using VectaMount (Vector Labs, H-5000).

### Clinical images

As part of the HARBOR clinical trial (NCT00891735)^13^, color fundus photographs, fluorescein angiography, and spectral-domain optical coherence tomography images (Cirrus; Carl Zeiss Meditec, Inc., Dublin, CA) were collected.

### Statistics for the *in vitro* analysis

Data are reported as the means ± standard deviation for the indicated number of experiments. At least three biological replicates were obtained for each experiment. Statistical analysis was carried out using the Prism v9 software. Statistical significance of continuous data was tested by the two-tailed Student’s t-test. p < 0.05 was considered statistically significant.

### Web Resources

dbSNP: https://www.ncbi.nlm.nih.gov/snp

Ensembl: http://grch37.ensembl.org/Homo_sapiens/Info/Index

OMIM: http://www.omim.org/

Uniprot: https://www.uniprot.org/

GTEx: https://www.gtexportal.org/home/

PolyPhen2: http://genetics.bwh.harvard.edu/pph2/

UK Biobank: https://www.ukbiobank.ac.uk/

Genebass: https://genebass.org/

## Data availability

All reagents used in this study are commercially available and supplier names/catalog numbers are provided in the Materials and Methods section of the manuscript. Human subjects were part of the HARBOR clinical trial, ClinicalTrials.gov identifier: NCT00891735, and the study population has been previously described for low-luminance deficit (LLD):

Frenkel, R.E., Shapiro, H., and Stoilov, I. (2016). Predicting vision gains with anti-VEGF therapy in neovascular age-related macular degeneration patients by using low-luminance vision. The British journal of ophthalmology 100, 1052-1057 http://doi.org/10.1136/bjophthalmol-2015-307575

Individual genetic data and other privacy-sensitive individual information are not publicly available because they contain information that could compromise research participant privacy. All publicly available code and software has been identified in the methods section of the manuscript. We are unable to share genome-wide individual level data, even de-identified, due to restrictions on the patient consents, however, all the summary statistics for the genetics analysis can be provided upon request to the corresponding author (Dr Marion Jeanne) and/or the lead Human Geneticist (Dr Brian Yaspan). Data is available for qualified researcher employed or legitimately affiliated with an academic, non-profit or government institution who have a track record in the field. We would ask the researcher to sign a data access agreement that needs to be signed by applicants and legal representatives of their institution, as well as legal representatives of Genentech, Inc. A brief research proposal will be needed to ensure that ‘Applications for access to Data must be Specific, Measurable, Attainable, Resourced and Timely.’ The following previously published datasets were used:

1. Human Retina and RPE/Choroid bulk RNA sequencing, data from: Orozco, L.D., Chen, H.H., Cox, C., Katschke, K.J., Jr., Arceo, R., Espiritu, C., Caplazi, P., Nghiem, S.S., Chen, Y.J., Modrusan, Z., et al. (2020). Integration of eQTL and a Single-Cell Atlas in the Human Eye Identifies Causal Genes for Age-Related Macular Degeneration. Cell Rep 30, 1246-1259 e1246. https://doi.org/10.1016/j.celrep.2019.12.082
2. Human eye single cell RNA sequencing, data from: Gautam, P., Hamashima, K., Chen, Y., Zeng, Y., Makovoz, B., Parikh, B.H., Lee, H.Y., Lau, K.A., Su, X., Wong, R.C.B., et al. (2021). Multi-species single-cell transcriptomic analysis of ocular compartment regulons. Nat Commun 12, 5675. https://doi.org/10.1038/s41467-021-25968-8

## Acknowledgments

We thank all of our Genentech colleagues involved in the Human Genetics Initiative including Julie Hunkapiller, Jens Reeder, and Suresh Selvaraj. We also thank our colleagues in the Research Pathology department, including Patrick Caplazi and Susan Haller.

## Competing interests

At the time of the study, all authors were full time employees of Genentech/Roche with stock and stock options in Roche. The funders had no role in study design, data collection and interpretation, or the decision to submit the work for publication.

## Supplemental files

Uploaded as 5 additional files:

1. Supplemental Figures S1 to S5 and Supplemental Table S1 (PDF file)
2. Supplemental Table S2 (xls file)
3. Source data file: Figure 6 – Source data 1 (PDF file) Uncropped scans of the films used to build figure 6 A, B, C and D. The area used in Figure 6 are highlighted on each film by red rectangles.
4. Reporting standards from the EQUATOR network: GRIPS checklist (PDF file)
5. MDAR checklist (PDF file)

## Rich media file

Figure 4 – Video 1: Example of a representative time-lapse video of a lipid droplet fusion event in pre-adipocytes expressing GFP-CIDEC wild-type (green) and mCherry-tagged PLIN1 (red).

## Abbreviations

AMD: Age-related macular degeneration
AR: autosomal recessive
BCVA: best corrected visual acuity
CIDEC: Cell-death-Inducing DNA fragmentation factor (DFF)45-like Effector C
CNV: choroidal neovascularization
EST: Expressed Sequence Tag
FPLD5: Familial Partial Lipodystrophy type 5
GA: Geographic Atrophy
GWAS: Genome-wide association studies
IAMDGC: International AMD Genetics Consortium
LD: lipid droplet
LLD: low-luminance deficit
LLVA: low-luminance visual acuity
MAF: minor allele frequency
MOI: mean optical intensity
OCT: Optical Coherence Tomography
OR: odds ratio
Q: quartile
RGCs: Retinal Ganglion Cells
RPE: Retinal Pigment Epithelium
SNP: single-nucleotide polymorphism
VEGF: Vascular Endothelial Growth Factor
WGS: whole genome sequencing
WT: wild-type

---

**Figure S1:**
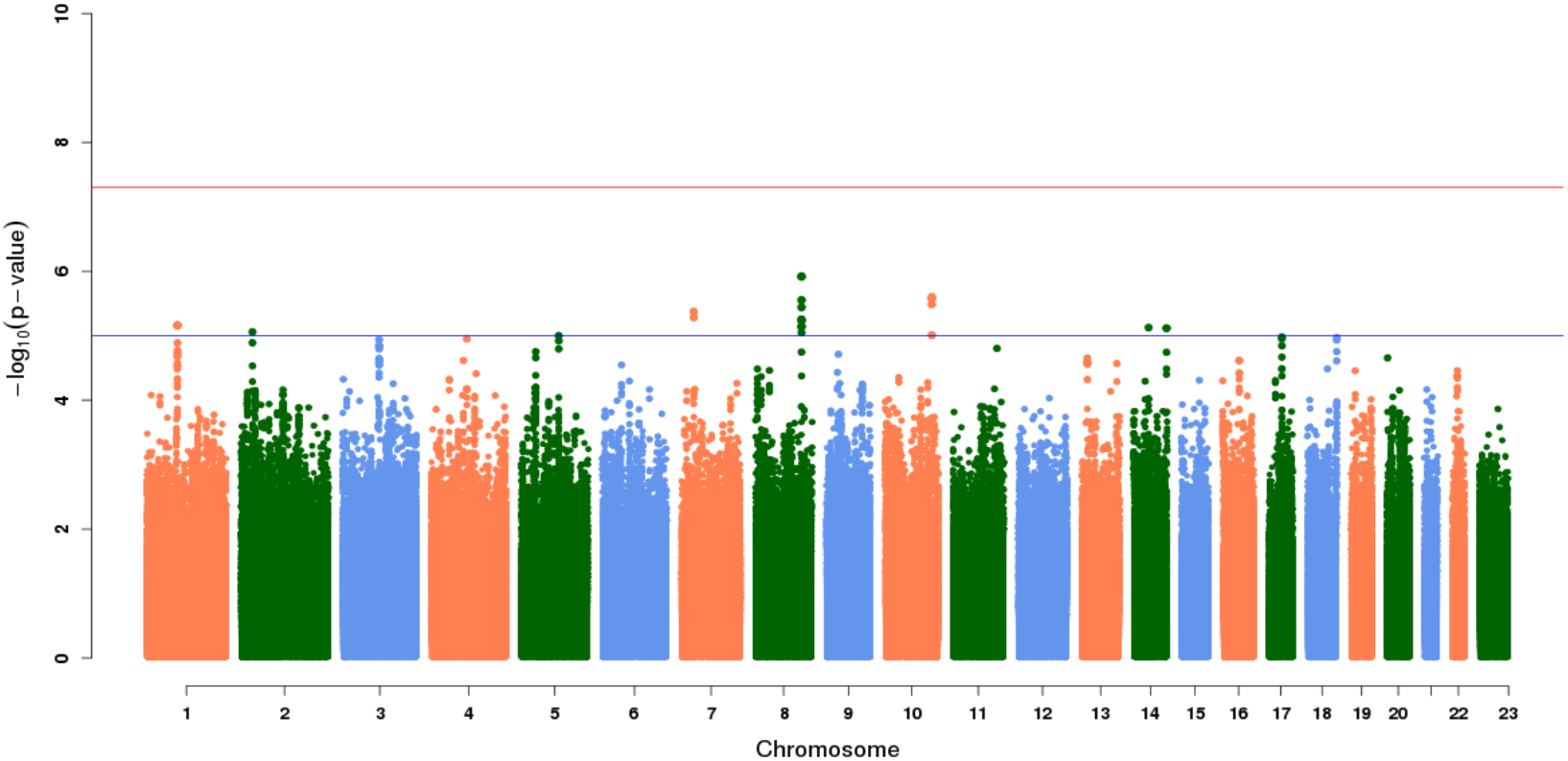
Manhattan plot of common variant analysis results contrasting AMD patients in the top and bottom LDD quartiles (Q1 and Q4).

**Figure S2:**
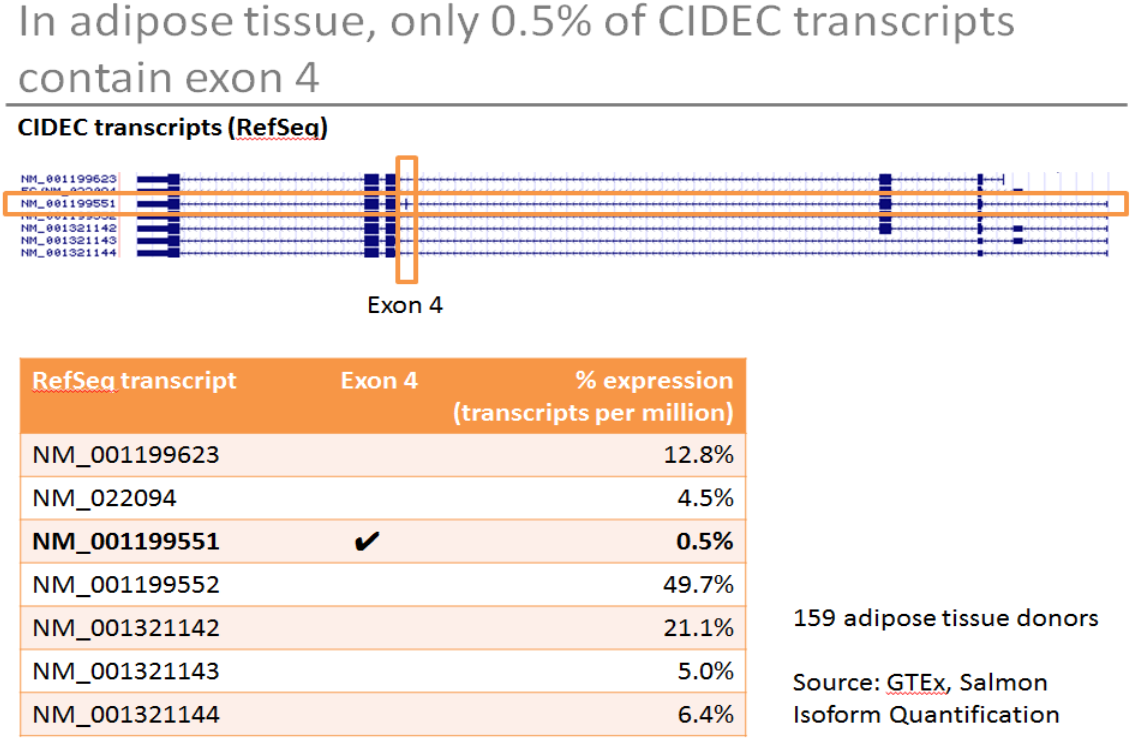
Summary of CIDEC exon expression in adipose tissue. *CIDEC* exon 4, the location of all rare variants seen in Q4 AMD patients, is expressed in 0.5% of all CIDEC transcripts found in adipose tissue in samples from the GTEx project.

**Figure S3:**
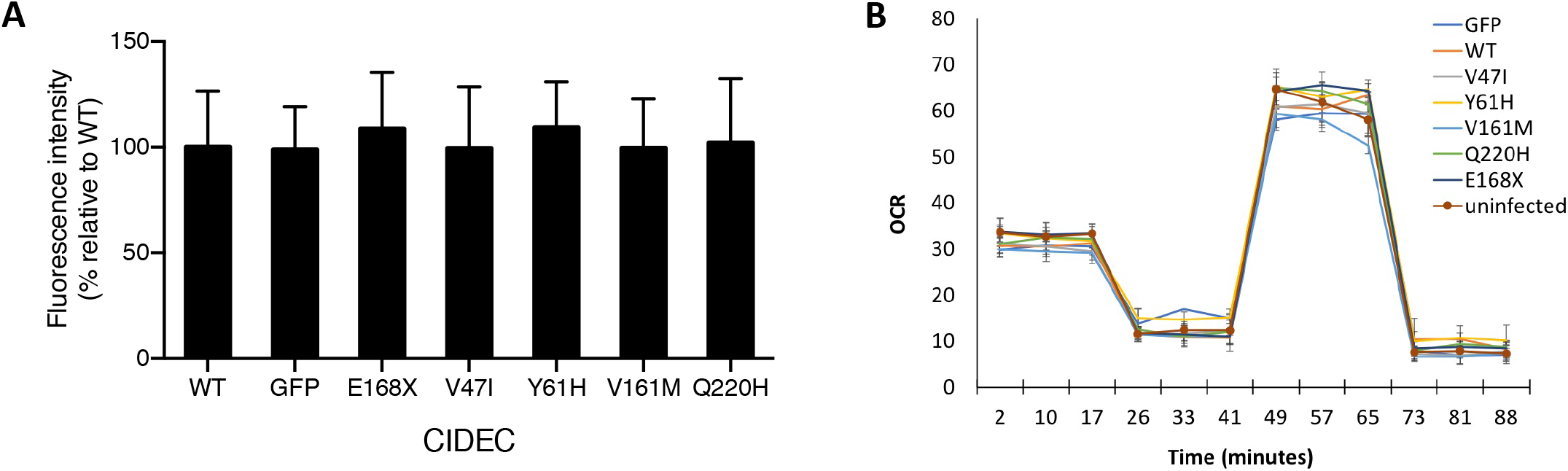
CIDEC rare variants do not affect mitochondria density or function. Quantification of mitochondria density using MitoTracker in 3T3-L1 cells expressing CIDEC wild-type (WT) or each of the rare variants (A). Mitochondria function measured by Seahorse analyzer (OCR: Oxygen consumption rate) (B).

**Figure S4:**
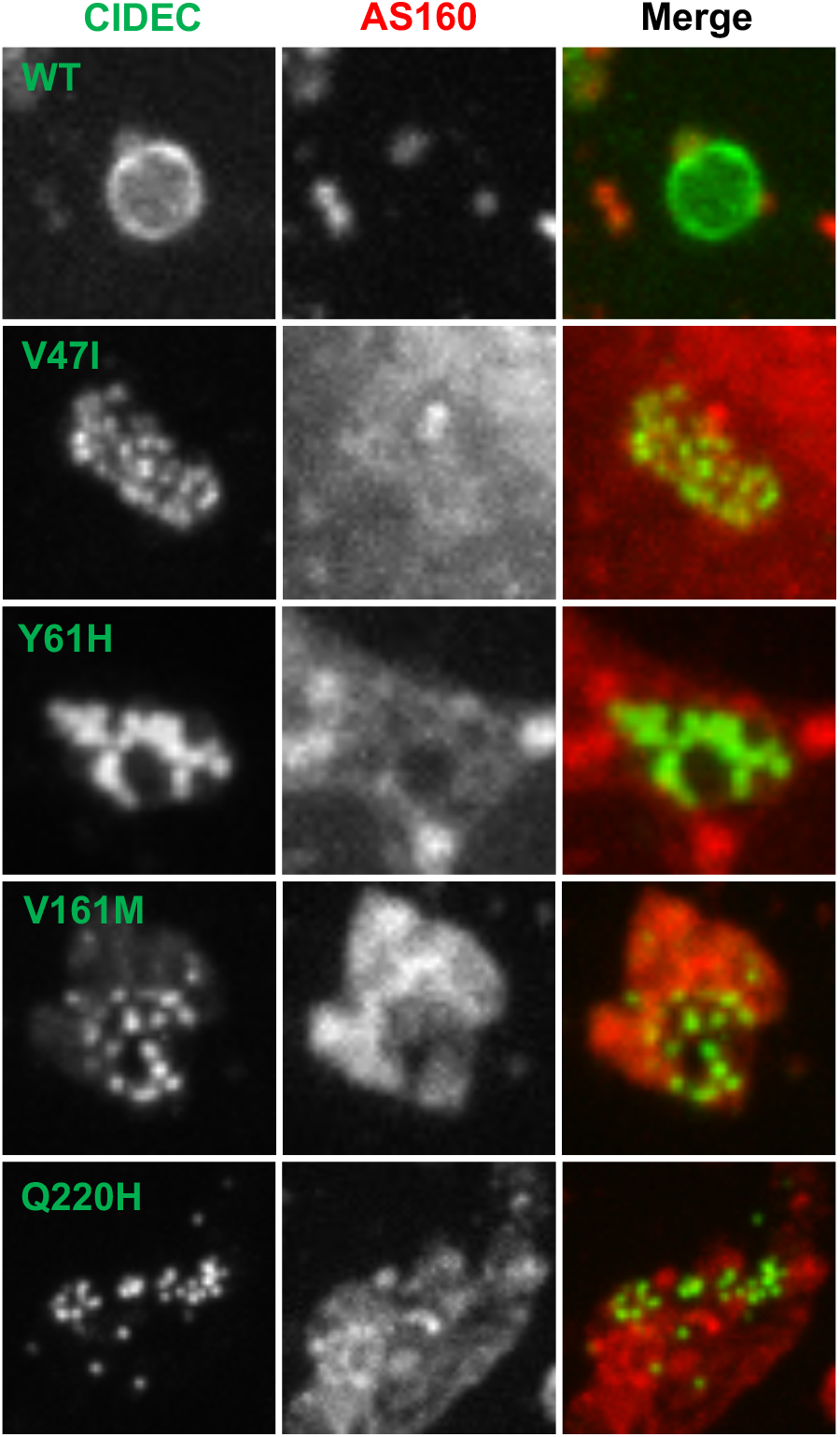
Representative images of CIDEC wild-type (WT) or rare variants (green) and AS160 (red) in pre-adipocytes.

**Figure S5:**
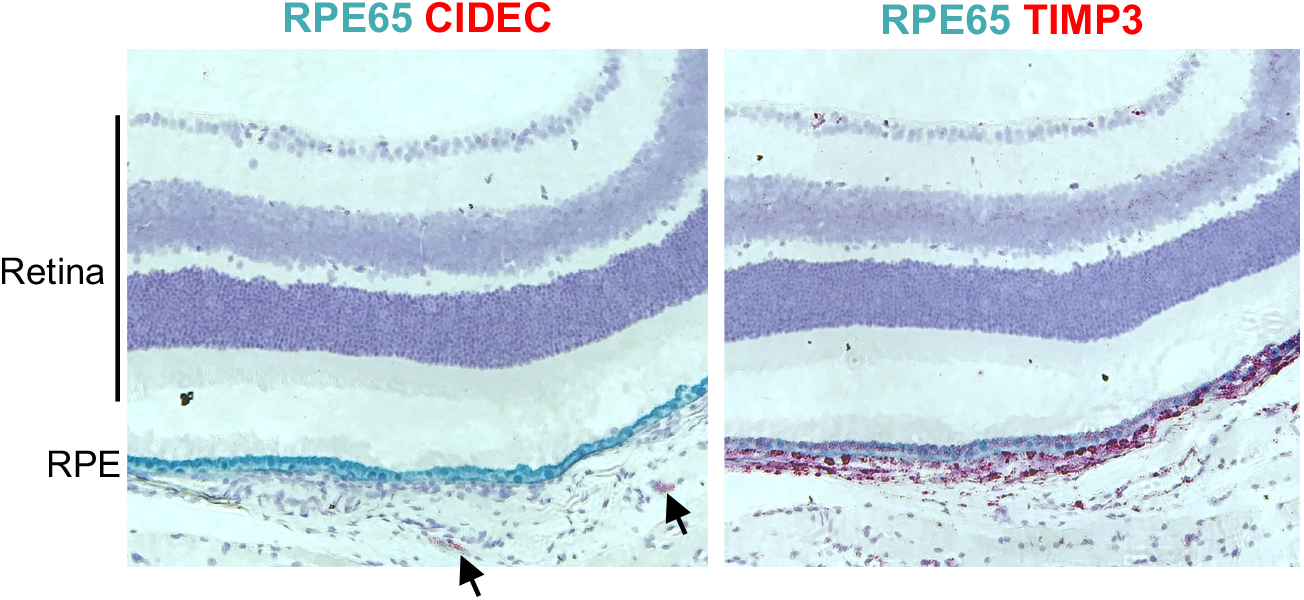
By in situ hybridization (ISH), Cidec RNA is not detected in mouse retina and Retinal Pigment Epithelium (RPE) cells. Rare Cidec positive cells are present in the choroidal tissue underneath the RPE (left: red, arrows). ISH for Rpe65 was used as RPE cell marker, and ISH for Timp3 (right: red) was used as positive control.

**Table S1:**
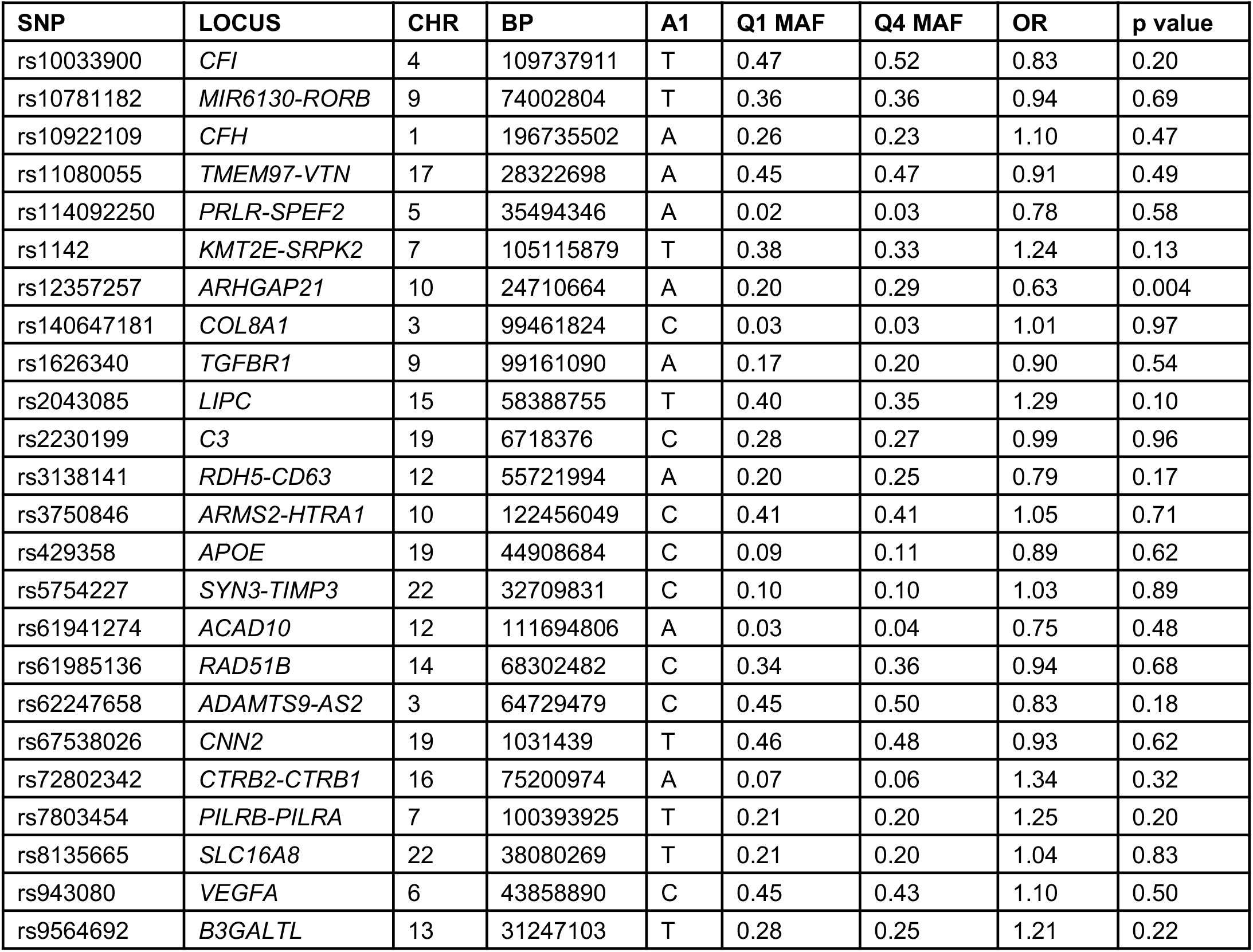
Comparison of AMD associated risk variants from Fritsche et. al, Nat Gen, 2015 in Q1 and Q4 AMD patients.

**Table S2:** (attached as a xls file) Results from UK Biobank rare variant burden PheWAS

## Notes

### Competing Interest Statement

All authors were employees of Genentech, Inc., a member of the Roche Group, and shareholder in Roche at the time of the study

## References

1. Wong, W.L., Su, X., Li, X., Cheung, C.M., Klein, R., Cheng, C.Y., and Wong, T.Y. (2014). Global prevalence of age-related macular degeneration and disease burden projection for 2020 and 2040: a systematic review and meta-analysis. Lancet Glob Health 2, e106–116.

2. Amoaku, W.M., Chakravarthy, U., Gale, R., Gavin, M., Ghanchi, F., Gibson, J., Harding, S., Johnston, R.L., Kelly, S.P., Lotery, A., et al. (2015). Defining response to anti-VEGF therapies in neovascular AMD. Eye (Lond) 29, 721–731.

3. Klein, M.L., Mauldin, W.M., and Stoumbos, V.D. (1994). Heredity and age-related macular degeneration. Observations in monozygotic twins. Archives of ophthalmology 112, 932–937.

4. Meyers, S.M., Greene, T., and Gutman, F.A. (1995). A twin study of age-related macular degeneration. American journal of ophthalmology 120, 757–766.

5. Hammond, C.J., Webster, A.R., Snieder, H., Bird, A.C., Gilbert, C.E., and Spector, T.D. (2002). Genetic influence on early age-related maculopathy: a twin study. Ophthalmology 109, 730–736.

6. Seddon, J.M., Ajani, U.A., and Mitchell, B.D. (1997). Familial aggregation of age-related maculopathy. American journal of ophthalmology 123, 199–206.

7. Heiba, I.M., Elston, R.C., Klein, B.E., and Klein, R. (1994). Sibling correlations and segregation analysis of age-related maculopathy: the Beaver Dam Eye Study. Genetic epidemiology 11, 51–67.

8. Seddon, J.M., Cote, J., Page, W.F., Aggen, S.H., and Neale, M.C. (2005). The US twin study of age-related macular degeneration: relative roles of genetic and environmental influences. Archives of ophthalmology 123, 321–327.

9. Fritsche, L.G., Igl, W., Bailey, J.N., Grassmann, F., Sengupta, S., Bragg-Gresham, J.L., Burdon, K.P., Hebbring, S.J., Wen, C., Gorski, M., et al. (2016). A large genome-wide association study of age-related macular degeneration highlights contributions of rare and common variants. Nat Genet 48, 134–143.

10. Sunness, J.S., Rubin, G.S., Applegate, C.A., Bressler, N.M., Marsh, M.J., Hawkins, B.S., and Haselwood, D. (1997). Visual function abnormalities and prognosis in eyes with age-related geographic atrophy of the macula and good visual acuity. Ophthalmology 104, 1677–1691.

11. Sunness, J.S., Rubin, G.S., Broman, A., Applegate, C.A., Bressler, N.M., and Hawkins, B.S. (2008). Low luminance visual dysfunction as a predictor of subsequent visual acuity loss from geographic atrophy in age-related macular degeneration. Ophthalmology 115, 1480–1488, 1488 e1481-1482.

12. Frenkel, R.E., Shapiro, H., and Stoilov, I. (2016). Predicting vision gains with anti-VEGF therapy in neovascular age-related macular degeneration patients by using low-luminance vision. The British journal of ophthalmology 100, 1052–1057.

13. Busbee, B.G., Ho, A.C., Brown, D.M., Heier, J.S., Suner, I.J., Li, Z., Rubio, R.G., Lai, P., and Group, H.S. (2013). Twelve-month efficacy and safety of 0.5 mg or 2.0 mg ranibizumab in patients with subfoveal neovascular age-related macular degeneration. Ophthalmology 120, 1046–1056.

14. Ho, A.C., Busbee, B.G., Regillo, C.D., Wieland, M.R., Van Everen, S.A., Li, Z., Rubio, R.G., Lai, P., and Group, H.S. (2014). Twenty-four-month efficacy and safety of 0.5 mg or 2.0 mg ranibizumab in patients with subfoveal neovascular age-related macular degeneration. Ophthalmology 121, 2181–2192.

15. Auer, P.L., and Lettre, G. (2015). Rare variant association studies: considerations, challenges and opportunities. Genome Med 7, 16.

16. Zhou, H., and Zhang, W. (2019). Gene expression profiling reveals candidate biomarkers and probable molecular mechanism in diabetic peripheral neuropathy. Diabetes Metab Syndr Obes 12, 1213–1223.

17. Karczewski, K.J., Solomonson, M., Chao, K.R., Goodrich, J.K., Tiao, G., Lu, W., Riley-Gillis, B.M., Tsai, E.A., Kim, H.I., Zheng, X., et al. (2021). Systematic single-variant and gene-based association testing of 3,700 phenotypes in 281,850 UK Biobank exomes. medRxiv, 2021.2006.2019.21259117.

18. Carithers, L.J., Ardlie, K., Barcus, M., Branton, P.A., Britton, A., Buia, S.A., Compton, C.C., DeLuca, D.S., Peter-Demchok, J., Gelfand, E.T., et al. (2015). A Novel Approach to High-Quality Postmortem Tissue Procurement: The GTEx Project. Biopreserv Biobank 13, 311–319.

19. Adzhubei, I.A., Schmidt, S., Peshkin, L., Ramensky, V.E., Gerasimova, A., Bork, P., Kondrashov, A.S., and Sunyaev, S.R. (2010). A method and server for predicting damaging missense mutations. Nat Methods 7, 248–249.

20. Liang, L., Zhao, M., Xu, Z., Yokoyama, K.K., and Li, T. (2003). Molecular cloning and characterization of CIDE-3, a novel member of the cell-death-inducing DNA-fragmentation-factor (DFF45)-like effector family. Biochem J 370, 195–203.

21. Chapman, A.B., Knight, D.M., Dieckmann, B.S., and Ringold, G.M. (1984). Analysis of gene expression during differentiation of adipogenic cells in culture and hormonal control of the developmental program. The Journal of biological chemistry 259, 15548–15555.

22. Danesch, U., Hoeck, W., and Ringold, G.M. (1992). Cloning and transcriptional regulation of a novel adipocyte-specific gene, FSP27. CAAT-enhancer-binding protein (C/EBP) and C/EBP-like proteins interact with sequences required for differentiation-dependent expression. The Journal of biological chemistry 267, 7185–7193.

23. Puri, V., Konda, S., Ranjit, S., Aouadi, M., Chawla, A., Chouinard, M., Chakladar, A., and Czech, M.P. (2007). Fat-specific protein 27, a novel lipid droplet protein that enhances triglyceride storage. The Journal of biological chemistry 282, 34213–34218.

24. Nishino, N., Tamori, Y., Tateya, S., Kawaguchi, T., Shibakusa, T., Mizunoya, W., Inoue, K., Kitazawa, R., Kitazawa, S., Matsuki, Y., et al. (2008). FSP27 contributes to efficient energy storage in murine white adipocytes by promoting the formation of unilocular lipid droplets. J Clin Invest 118, 2808–2821.

25. Toh, S.Y., Gong, J., Du, G., Li, J.Z., Yang, S., Ye, J., Yao, H., Zhang, Y., Xue, B., Li, Q., et al. (2008). Up-regulation of mitochondrial activity and acquirement of brown adipose tissue-like property in the white adipose tissue of fsp27 deficient mice. PLoS One 3, e2890.

26. Rubio-Cabezas, O., Puri, V., Murano, I., Saudek, V., Semple, R.K., Dash, S., Hyden, C.S., Bottomley, W., Vigouroux, C., Magre, J., et al. (2009). Partial lipodystrophy and insulin resistant diabetes in a patient with a homozygous nonsense mutation in CIDEC. EMBO Mol Med 1, 280–287.

27. Zhou, L., Park, S.Y., Xu, L., Xia, X., Ye, J., Su, L., Jeong, K.H., Hur, J.H., Oh, H., Tamori, Y., et al. (2015). Insulin resistance and white adipose tissue inflammation are uncoupled in energetically challenged Fsp27-deficient mice. Nat Commun 6, 5949.

28. Sun, Z., Gong, J., Wu, H., Xu, W., Wu, L., Xu, D., Gao, J., Wu, J.W., Yang, H., Yang, M., et al. (2013). Perilipin1 promotes unilocular lipid droplet formation through the activation of Fsp27 in adipocytes. Nat Commun 4, 1594.

29. Walther, T.C., and Farese, R.V., Jr. (2009). The life of lipid droplets. Biochim Biophys Acta 1791, 459–466.

30. Bersuker, K., Peterson, C.W.H., To, M., Sahl, S.J., Savikhin, V., Grossman, E.A., Nomura, D.K., and Olzmann, J.A. (2018). A Proximity Labeling Strategy Provides Insights into the Composition and Dynamics of Lipid Droplet Proteomes. Dev Cell 44, 97–112 e117.

31. Gao, G., Chen, F.J., Zhou, L., Su, L., Xu, D., Xu, L., and Li, P. (2017). Control of lipid droplet fusion and growth by CIDE family proteins. Biochim Biophys Acta Mol Cell Biol Lipids 1862, 1197–1204.

32. Wu, C., Zhang, Y., Sun, Z., and Li, P. (2008). Molecular evolution of Cide family proteins: novel domain formation in early vertebrates and the subsequent divergence. BMC Evol Biol 8, 159.

33. Wu, L., Xu, D., Zhou, L., Xie, B., Yu, L., Yang, H., Huang, L., Ye, J., Deng, H., Yuan, Y.A., et al. (2014). Rab8a-AS160-MSS4 regulatory circuit controls lipid droplet fusion and growth. Dev Cell 30, 378–393.

34. Orozco, L.D., Chen, H.H., Cox, C., Katschke, K.J., Jr., Arceo, R., Espiritu, C., Caplazi, P., Nghiem, S.S., Chen, Y.J., Modrusan, Z., et al. (2020). Integration of eQTL and a Single-Cell Atlas in the Human Eye Identifies Causal Genes for Age-Related Macular Degeneration. Cell Rep 30, 1246–1259 e1246.

35. Gautam, P., Hamashima, K., Chen, Y., Zeng, Y., Makovoz, B., Parikh, B.H., Lee, H.Y., Lau, K.A., Su, X., Wong, R.C.B., et al. (2021). Multi-species single-cell transcriptomic analysis of ocular compartment regulons. Nat Commun 12, 5675.

36. Imanishi, Y., Batten, M.L., Piston, D.W., Baehr, W., and Palczewski, K. (2004). Noninvasive two-photon imaging reveals retinyl ester storage structures in the eye. J Cell Biol 164, 373–383.

37. McBee, J.K., Palczewski, K., Baehr, W., and Pepperberg, D.R. (2001). Confronting complexity: the interlink of phototransduction and retinoid metabolism in the vertebrate retina. Prog Retin Eye Res 20, 469–529.

38. Song, F.Q., Zhou, H.M., Ma, W.X., Li, Y.L., Hu, B.A., Shang, Y.Y., Wang, Z.H., Zhong, M., Zhang, W., and Ti, Y. (2022). CIDEC: A Potential Factor in Diabetic Vascular Inflammation. J Vasc Res 59, 114–123.

39. Rajamoorthi, A., Lee, R.G., and Baldan, A. (2018). Therapeutic silencing of FSP27 reduces the progression of atherosclerosis in Ldlr(-/-) mice. Atherosclerosis 275, 43–49.

40. van Leeuwen, E.M., Emri, E., Merle, B.M.J., Colijn, J.M., Kersten, E., Cougnard-Gregoire, A., Dammeier, S., Meester-Smoor, M., Pool, F.M., de Jong, E.K., et al. (2018). A new perspective on lipid research in age-related macular degeneration. Prog Retin Eye Res 67, 56–86.

41. Tanaka, N., Takahashi, S., Matsubara, T., Jiang, C., Sakamoto, W., Chanturiya, T., Teng, R., Gavrilova, O., and Gonzalez, F.J. (2015). Adipocyte-specific disruption of fat-specific protein 27 causes hepatosteatosis and insulin resistance in high-fat diet-fed mice. The Journal of biological chemistry 290, 3092–3105.

42. Zhou, L., Yu, M., Arshad, M., Wang, W., Lu, Y., Gong, J., Gu, Y., Li, P., and Xu, L. (2018). Coordination Among Lipid Droplets, Peroxisomes, and Mitochondria Regulates Energy Expenditure Through the CIDE-ATGL-PPARalpha Pathway in Adipocytes. Diabetes 67, 1935–1948.

43. Lyzogubov, V.V., Tytarenko, R.G., Bora, N.S., and Bora, P.S. (2012). Inhibitory role of adiponectin peptide I on rat choroidal neovascularization. Biochim Biophys Acta 1823, 1264–1272.

44. Fu, Z., Liegl, R., Wang, Z., Gong, Y., Liu, C.H., Sun, Y., Cakir, B., Burnim, S.B., Meng, S.S., Lofqvist, C., et al. (2017). Adiponectin Mediates Dietary Omega-3 Long-Chain Polyunsaturated Fatty Acid Protection Against Choroidal Neovascularization in Mice. Investigative ophthalmology & visual science 58, 3862–3870.

45. Li, H.Y., Hong, X., Cao, Q.Q., and So, K.F. (2019). Adiponectin, exercise and eye diseases. Int Rev Neurobiol 147, 281–294.

46. Cao, G., Chen, Y., Zhang, J., Liu, Y., Zhang, M., Zhang, K., and Su, Z. (2015). Effects of adiponectin polymorphisms on the risk of advanced age-related macular degeneration. Biomarkers 20, 266–270.

47. Kaarniranta, K., Paananen, J., Nevalainen, T., Sorri, I., Seitsonen, S., Immonen, I., Salminen, A., Pulkkinen, L., and Uusitupa, M. (2012). Adiponectin receptor 1 gene (ADIPOR1) variant is associated with advanced age-related macular degeneration in Finnish population. Neurosci Lett 513, 233–237.

